# Improved inference of multiscale sequence statistics in generative protein models

**DOI:** 10.64898/2026.04.06.716859

**Authors:** Marion Chauveau, Yaakov Kleeorin, Emily Hinds, Ivan Junier, Rama Ranganathan, Olivier Rivoire

## Abstract

High dimensionality and multiscale statistical structure are pervasive features of biological data, posing fundamental challenges for modeling. Because model inference generally proceeds with far fewer data than parameters, statistical patterns across scales are often unevenly represented. Protein sequences provide a paradigmatic example: statistics across homologs are inherently multiscale, displaying collective correlations among conserved residue sectors that encode function, alongside localized correlations corresponding to physical contacts outside these sectors. Standard regularization strategies used to mitigate undersampling during model inference have been shown to capture these patterns unevenly, a bias that compromises generative models of protein sequences by limiting their ability to produce both functional and diverse proteins. This limitation is exemplified by Boltzmann Machine–based generative models, which so far have required post hoc corrections to recover functionality, at the cost of reduced sequence diversity and novelty. Here, we introduce the stochastic Boltzmann Machine (sBM), a new regularization strategy that more accurately captures different correlation scales. Through analyses of theoretical models with known ground-truth parameters and experiments on the chorismate mutase family, we show that sBM effectively mitigates distortions in the estimation of model parameters, enabling the generation of functional sequences with greater diversity and without the need for post hoc corrections. These results advance the inference of generative models that more faithfully reflect the evolutionary constraints shaping protein sequences.

## Introduction

A central challenge in biology is to describe high-dimensional systems of interacting components that exhibit correlations across multiple scales, from collective modes involving many variables to localized interactions among few. Protein sequences exemplify this challenge: analyses of conservation and correlations in multiple sequence alignments of homologous proteins reveal a rich multiscale statistical structure. Quantitatively characterizing these patterns has motivated a broad range of approaches, spanning correlation-based methods [1, 2], maximum-entropy models [3, 4], and modern machine-learning techniques [5]. These approaches have yielded important results, from predicting residue-residue contacts [3, 6, 7] and allosteric pathways [1, 8] to estimating mutational effects [9] and designing functional proteins [10, 11]. Among these approaches, generative models are particularly compelling, as they not only capture the statistical structure of natural sequences but also enable the generation of synthetic ones [12]. Early demonstrations were provided by statistical coupling analysis (SCA) [10], followed by models inspired by statistical physics and machine learning, including Potts models [11], variational autoencoders [13], diffusion models [14], and transformers [15, 16]. Despite differences in their architectures and levels of complexity, all of these models face similar conceptual and practical challenges.

A first challenge, common to all generative models, is the need to evaluate them across multiple dimensions. Previous work on inferring models from multiple sequence alignments has emphasized reproducing empirical statistics or predicting structural contacts [4, 13, 15, 17, 18], but optimally reproducing or predicting specific properties of the data does not guarantee optimal generativity. Instead, three complementary criteria are widely recognized as essential for assessing generative performance [19, 20, 21]: (1) fidelity, here the extent to which the generated sequences have functional properties comparable to natural ones; (2) novelty, the extent to which these sequences are distinct from the training sequences; (3) diversity, the extent to which they span the full range of natural sequence variation (to be distinguished from entropy, which quantifies the total span without reference to natural variation). For proteins, novelty and diversity can be assessed from sequence analyses, whereas fidelity can be tested experimentally. Such experimental evaluation has so far been carried out for only a few protein families, limiting systematic comparisons between different generative models [10, 11, 22, 13, 23, 14, 24]. Nevertheless, available data indicate that one of the simplest models, Potts model, achieve state-of-the-art performance. This has been demonstrated in the context of the AroQ family of chorismate mutases [11], enzymes that catalyze a key step in the biosynthesis of tyrosine and phenylalanine, and whose activity can be assayed in vivo, which makes them an ideal experimental test case.

A second challenge for generative models stems from limited and uneven sampling. In the context of proteins, limited sampling arises because the number of available homologous sequences is typically insufficient to reliably determine all the model parameters. Uneven sampling arises from shared evolutionary histories, which implies the non-independence of these sequences, and from biased representation in protein databases. Here, we follow standard practice and use sequence reweighting to mitigate uneven sampling at a local scale. Beyond this correction, we strive to capture all statistically significant patterns, regardless of their functional or phylogenetic origins, which are often intertwined. Our focus is instead on overcoming the limitations imposed by limited sampling.

A common strategy to mitigate undersampling effects is the introduction of regularization, which constrains the range of admissible parameter values. In Potts models, this is typically achieved using *L*_2_ regularization, which penalizes all parameters uniformly [4]. However, such a uniform regularization introduces systematic biases when applied to data exhibiting statistical patterns of different scales, as is the case for protein sequences [25]. Sequence alignments exhibit two such statistical patterns [26, 27]: collective correlations among multiple residues (sectors) reflecting functional constraints related to binding, catalysis, or allostery [28], and localized correlations outside of these sectors reflecting stability constraints on some contacting residues [29]. Because uniform regularization captures these two correlation scales unevenly, sequences sampled from generative models using such regularization are predominantly nonfunctional [11]. This negative outcome can be partially mitigated through rescaling of model parameters (corresponding to introducing a “temperature” parameter and sampling at low temperature) at the expense of diminished diversity and novelty [11]. However, in the presence of an experimental bias, the effectiveness of this approach depends critically on the incidental alignment between this bias and the sampling bias introduced by the post hoc correction (Fig. S1). More fundamentally, it does not address the challenge of modeling all relevant constraints evenly.

In this work, we introduce a novel regularization strategy for inferring Potts model. In contrast to the previous Boltzmann Machine (BM) [30] inference method, our new approach, termed stochastic Boltzmann Machine (sBM), reduces systematic biases and generates functional, diverse protein sequences without requiring post hoc rescaling, as we demonstrate both in the context of mathematically defined models and in the context of experiments with the chorismate mutase enzyme family. Although applied here to proteins within a specific inference framework, our approach may have broader relevance for model inference in biological systems exhibiting multiple statistical patterns.

## Methods

### Potts models

Potts models are inferred from a multiple sequence alignment (MSA) of *M* homologous proteins, denoted as ***σ*** = (*σ*_1_*, σ*_2_*, …, σ_L_*), where *σ_i_* represents one of *q* different amino acids, and *L* is the length of the sequences. Potts models are probabilistic models where the distribution has a Boltzmann form *P* (***σ***) = exp (−*E*(***σ***)) */Z*(***h***, ***J*** ) with an “energy” given by

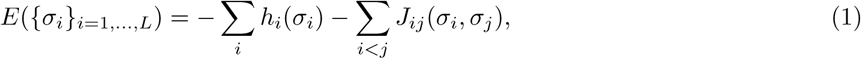

where *Z*(***h***, ***J*** ) is a normalizing constant, referred to as the partition function. The *h_i_*(*σ_i_*)’s and the *J_ij_*(*σ_i_, σ_j_*)’s are the model parameters respectively called the fields and the couplings. They are chosen to maximize the log-likelihood L(***h***, ***J*** ) = (1*/M* )Σ***_σ_*** ln *P* (***σ***|***h***, ***J*** ) where the sum is over the *M* sequences of the alignment. This maximum is achieved when the single and pairwise frequencies *f_i_*(*a*) and *f_ij_*(*a, b*) of amino acids *a, b* at individual positions *i* and pairs of positions *i, j* are matched between the model and the data. This makes transparent what features of the data are incorporated in the model but also why a naïve optimization is prone to overfitting: the statistics from the alignment are indeed subject to sampling noise due to the limiting number *M* of sequences, and reproducing exactly those statistics is generally not desirable.

### Boltzmann Machine (BM) inference

To reduce the impact of undersampling and improve model robustness, it is common to introduce regularization that limits the effective amplitude of the inferred parameters. A typical implementation is *L*_2_ regularization, which penalizes large parameter values through a quadratic constraint on the fields and couplings [31, 9, 32, 4, 11, 25, 33]. The optimization is then performed on L(***h***, ***J*** ) − *λ_h_*Σ*_i_*Σ*_a_h_i_*(*a*)^2^ − *λ_J_*Σ*_i<j_*Σ*_a,b_ J_ij_*(*a, b*)^2^ where *λ_h_* and *λ_J_* control the regularization strength for fields and couplings, respectively (see Supplementary Methods).

Once the objective function is defined, the model parameters are typically inferred using vanilla gradient descent, an iterative optimization procedure that, for the couplings, takes the form

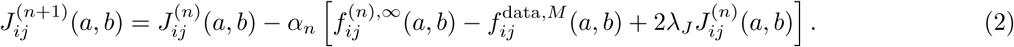

and similarly for the fields (see Supplementary Methods). Here, 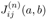 denotes the estimate of *J_ij_*(*a, b*) at iteration 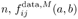 is the empirical frequency of (*a, b*) at sites (*i, j*) in the *M* sequences of the dataset, 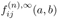 is the corresponding frequency estimated from the model parameters at iteration *n*, and *α_n_* is the learning rate. Following standard practice in the field to partly correct for the uneven sampling of sequence space, empirical frequencies are obtained by weighting the contribution of each sequence *s* with a weight *w_s_* inversely proportional to the number of sequences similar to *s*, thus defining an effective number of sequences 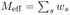 lower than the actual number of sequences *M* (see Supplementary Methods). The model frequencies 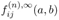 are generally prohibitively expensive to compute exactly and are typically approximated by Monte Carlo sampling using a sufficiently large number of samples (indicated by the symbol ∞) to perform the average (see Supplementary Methods). Following earlier work [30], we refer to this inference method as the Boltzmann Machine (BM).

### Stochastic Boltzmann Machine (sBM) inference

To address the limitations of BM inference, we introduce the stochastic Boltzmann Machine (sBM) which differs from vanilla gradient descent in two ways.

First, it leverages not only gradient information but also a local approximation of the curvature of the objective function, which reveals which parameter directions are stiff (where small changes have a major impact) and which are sloppy (where large changes have minimal effect). This alone is not novel: quasi-Newton methods incorporating curvature information have previously accelerated BM inference [34, 35]. Although our method also converges substantially faster than standard BM, our primary motivation for adopting a second-order approach is its ability to better handle anisotropic curvature of the log-likelihood in parameter space, as induced by the rich statistical structure of the data.

Second, the sBM replaces explicit and uniform regularization, such as *L*_2_ penalties, with implicit regularization that emerges from the inference process itself. This implicit regularization combines three complementary mechanisms, each controlled by a distinct hyperparameter: (i) *early stopping*, with a limited number *N*_iter_ of iterations in gradient descent [36]; (ii) *approximate curvature estimation* through the Limited-memory Broyden-Fletcher-Goldfarb-Shanno (L-BFGS) algorithm [37], with a parameter *m* controlling the amount of curvature information retained from previous iterations, and therefore the fidelity of the approximation; (iii) *limited sampling* of model statistics using a finite number *N*_chains_ of Monte Carlo samples, as a way to reflect the limited number *M* of sequences to estimate statistics from the data. As we will show, this novel regularization treats statistical patterns across scales more evenly, thereby reducing the biases observed with explicit and uniform regularization.

Accordingly, Eq. (2) becomes

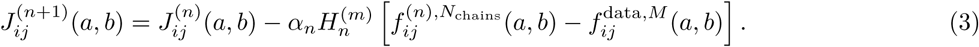

where 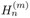 denotes the Hessian approximation restricted to the coupling parameters, and *f* ^(^*^n^*^)^*^,N^*^chains^ is the model-estimated frequency at iteration *n*, computed from *N*_chains_ independent Monte Carlo chains. The total number of iterations *N*_iter_, not shown explicitly in the equation, determines the value of *n* at which optimization is stopped. We refer to this inference method as the sBM, by contrast with the standard BM that does not involve the Hessian and takes the limits *N*_chains_ → ∞ and *N*_iter_ → ∞. In practice, *N*_chains_ is usually taken to be of the order of the number of effective sequences in the alignment (*M*_eff_), while the number of iterations *N*_iter_ is stopped as soon as the empirical statistics 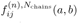 match the empirical statistics 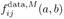 within some predefined accuracy, e.g., as soon as the Pearson correlation coefficient between the inferred and empirical connected correlations *C_ij_*(*a, b*) = *f_ij_*(*a, b*) − *f_i_*(*a*)*f_j_*(*b*) reaches a threshold of 0.95 [38].

Here we adopt different criteria to select *N*_iter_ and *N*_chains_, which lead to very different (smaller) values of these parameters. We use a small *N*_chains_ to enforce the computation of model statistics in an undersampled regime and choose *N*_iter_ based on features of the iterative dynamics. In these dynamics, the total norm of the inferred parameters diverges at a non-constant rate, with a plateau at an intermediate number of iterations from which *N*_iter_ is chosen (Fig. S2). A small *m* helps slow down the dynamics and extend this plateau, facilitating this choice (Fig. S3). Setting *m* = 1, this procedure leaves essentially only one hyperparameter, *N*_chains_, to be set. The value of *N*_chains_ achieving the optimal trade-off between fidelity, novelty, and diversity requires experiments to measure fidelity, but a range of values can be predetermined without experiments by selecting values of *N*_chains_ with comparable levels of novelty to the original data (Fig. S4). The three components of implicit regularization have complementary effects that reduce overfitting, and each is necessary for the effects we present (Fig. S3). In particular, approximate curvature estimation via the L-BFGS algorithm is necessary beyond its role in accelerating convergence (Fig. S2). Finally, as a small value of *N*_chains_ introduces variability across independent runs (Fig. S5), we average the inferred fields and couplings over multiple independent runs (10 by default and 50 for the models selected for experimental characterization) to obtain reproducible models. In contrast to other inference methods, we reinitialize the Monte Carlo sampling at each iteration step and do not reuse the state from previous steps, as in persistent contrastive divergence implemented in other approaches [38, 39], which leads to qualitatively different results (Fig. S6). Also in contrast to other inference methods, we do not use pseudocounts to modify empirical statistics, which has a different effect from that of our regularization parameters (Fig. S11).

## Results

Following the evaluation procedures established in previous studies [25, 11], we first compare how the BM and sBM differ in the biases they introduce through the interplay between undersampling and statistical structure. We then assess their ability to generate functional, novel, and diverse sequences. These comparisons are performed in two settings: first, using synthetic data generated from a mathematically defined teacher model illustrated in Fig. 1 (see also Supplementary Methods), and second, using natural protein data from the chorismate mutase family. Each setting presents complementary advantages and limitations. Synthetic data allow for direct and quantitative validation by enabling a comparison between inferred and teacher parameters, which is not possible with real sequences. However, this approach depends on the choice of the teacher model, which must adequately capture the main features of natural sequence statistics. Here we adopt the highly simplified teacher model introduced in [25], designed to contain two clearly separated scales. This model takes the form of Eq. (1), where localized interactions correspond to isolated pairwise couplings, and collective interactions are modeled as collective units composed of pairwise couplings (Fig. 1A). While other studies have used as benchmarks teacher models inferred from multiple sequence alignments of actual proteins [40, 41, 42], we deliberately avoid this approach here, as our point is to correct biases that these inferred models typically present. Instead, to demonstrate that our results apply to real proteins, we directly compare our analysis of an alignment of the chorismate mutase with new experimental results. This approach avoids modeling assumptions, though it is constrained by experimental limitations such as the number of sequences that we can test and intrinsic experimental biases. Despite the respective limitations of synthetic and real data, obtaining consistent results in both contexts provides an effective form of cross-validation.

**Figure 1:**
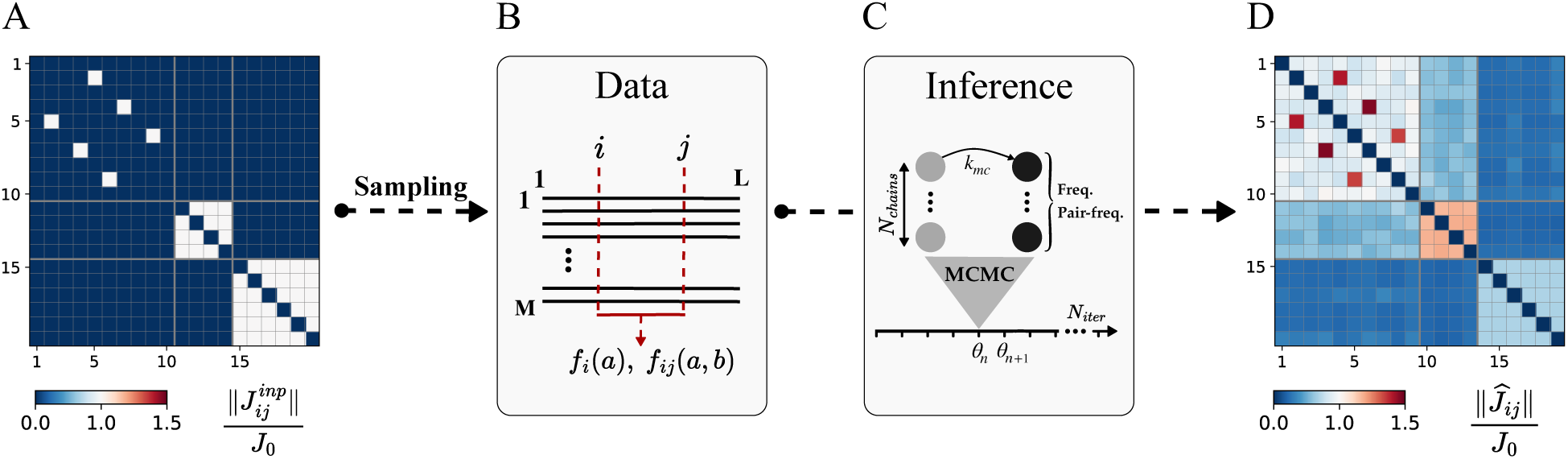
Inference with synthetic data from a mathematical model. The model proposed by Kleeorin *et al.* [25] aims to capture a fundamental characteristic of proteins: the presence of interaction between residues at various scales. It takes the form of Eq. (1) with *L* = 20 positions, *q* = 10 possible amino acids at each site, and no fields *h_i_*. **A**. The *L* × *L* Frobenius norm of the couplings *J* ^inp^(*a, b*) of the teacher model, with three types of features: isolated pairwise couplings at positions (2,5), (4,7) and (6,9), a small collective unit at positions (11-14) and a larger collective unit at positions (15-20). The couplings are normalized by the Frobenius value in the zero-sum gauge *J*_0_. **B**. To reproduce the undersampling observed in natural protein families, *M* = 300 sequences are sampled from the teacher model, and their single-site and pairwise frequencies are computed. These data are then used to infer the model parameters. **C**. Schematic illustration of the model inference with a gradient descent algorithm. The model parameters at iteration *n* of the gradient descent are stored in the vector ***θ****n*. At each step, a Monte Carlo simulation is used to sample *N*_chains_ sequences from the model. The sequences obtained after *k*_mc_ MCMC steps are then used to compute single-site and pairwise frequencies, which are required to evaluate the gradient of the objective function. **D**. The *L* × *L* Frobenius norm of the inferred couplings *Ĵ_ij_* (*a, b*) in the zero-sum gauge, ∥*Ĵ_ij_* ∥, normalized by *J*_0_, using the Boltzmann Machine with *L*_2_ regularization hyperparameter *λ_J_* = 10*^−^*^3^. The comparison with A shows biases induced by the uniform regularization: couplings associated with the collective pattern are underestimated while couplings associated with isolated pairs are overestimated.

### Inferred biases in the context of synthetic data

Previous results obtained with the BM inference and explicit (*L*_2_) regularization on synthetic data generated from the teacher model of Fig. 1 and Fig. 2A are summarized in Fig. 2B. Three main observations emerge [25]: (1) biases in inferred couplings, with isolated interactions being overestimated at low regularization and underestimated at high regularization relative to collective ones; (2) a strong sensitivity of these biases to the choice of regularization strength; (3) the absence of any single value of the regularization hyperparameter that allows all coupling types to be inferred accurately.

**Figure 2:**
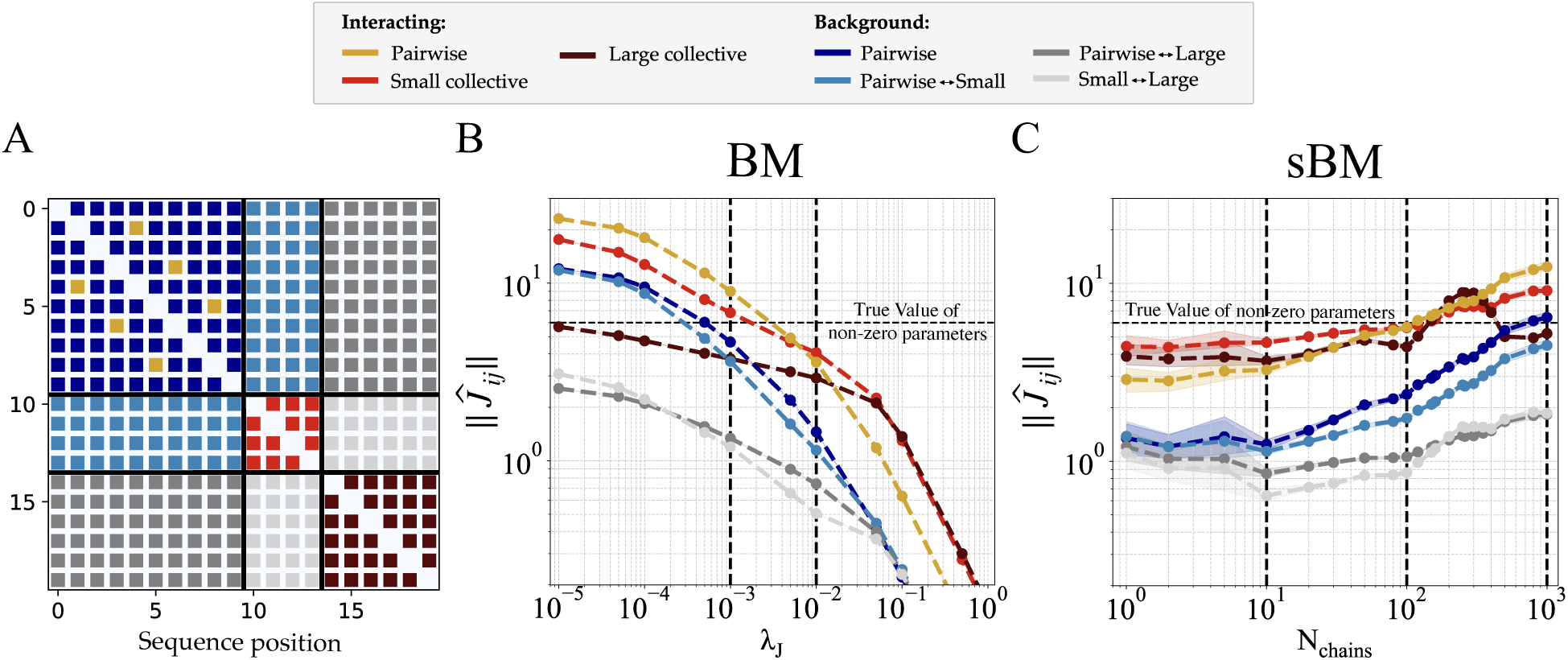
BM and sBM inference in the context of synthetic data. The synthetic dataset consists of *M* = 300 sequences generated from the mathematical model presented in Fig. 1. **A**. *L*×*L* matrix of the teacher model, showing the same interaction structure as in Fig. 1A: isolated pairwise couplings at positions (2,5), (4,7), and (6,9) (yellow); a small collective unit spanning positions 11-14 (light red); and a large collective unit at positions 15-20 (dark red). Non-interacting positions are divided into four groups corresponding to the background of pairwise couplings (dark blue); the background between pairwise couplings and the small collective unit (light blue); the background between pairwise couplings and the large collective unit (grey); and the background between the two collective units (light grey). **B**. BM inference. The inference is done using vanilla gradient descent, with parameters *N*_chains_ = 1000, *k*_mc_ = 10^5^, *N*_iter_ = 3 · 10^4^, *L*_2_ regularization with *λ_h_*= 0.01, and *λ_J_*as a tunable parameter. We show the magnitude of inferred couplings as a function of *λ_J_* . In the low-regularization regime, isolated contacts dominate, while strong regularization emphasizes collective features. The vertical dotted lines indicate the regularization levels used in Fig. 4. **C**. sBM inference. The inference is done with the L-BFGS algorithm with *m* = 1, *k*_mc_ = 10^5^, *N*_iter_ = 400. *N*_chains_ is varied to control the strength of regularization. We show the magnitude of inferred couplings as a function of *N*_chains_ (with large *N*_chains_ corresponding to weak regularization). A wide range of *N*_chains_ yields nearly unbiased estimates of interacting couplings. The vertical dotted lines mark the regularization level used in Fig. 4. In panels B and C, each point is the mean of 10 independent repeats. BM: average over 10 independently inferred BMs. sBM: average over 10 sBMs, where the parameters of each sBM are themselves the average of 10 independently inferred models. Error bars represent standard deviation obtained across the 10 repeats.

We repeated the analysis using the sBM on the same synthetic dataset. Fig. 2C shows the inferred couplings as a function of the main regularization parameter, *N*_chains_, with *m* = 1 and *N*_iter_ = 400 (see Fig. S7 for variations in these parameters, and Fig. S8 for a more direct comparison to ground-truth values). For large *N*_chains_ ∼ 1000, the sBM reproduces the same biases as the BM under weak *L*_2_ regularization, namely an overestimation of isolated pairwise couplings and small collective interactions. At very low *N*_chains_, the bias reverses: isolated couplings become slightly weaker than collective ones, and we observe increased variability across inference runs, even after averaging over ten models as done here. However, unlike the BM inference, increasing the regularization in the sBM (that is, decreasing *N*_chains_) does not substantially dampen the overall coupling magnitudes. More importantly, there exists a range of intermediate *N*_chains_ values for which interacting couplings are inferred with limited bias and remain close to their true values, in contrast to what is obtained with BM. Finally, although non-interacting couplings are not driven to zero, their magnitudes remain systematically smaller than those of true interactions.

In the context of our synthetic data, sBM thus resolves the biases of the standard BM: appropriate hyperparameters exist that enable patterns of different scales to be inferred with correct relative values.

### Biases in the context of protein data

In previous work, we showed that similar biases arise when applying the BM to a multiple sequence alignment of the chorismate mutase family [25]. Although no ground-truth model exists for this system, evolutionary analyses and experiments probing mutational effects distinguish local from collective correlation patterns (Fig.3A): both are evident in the MSA as residue correlations, but collective patterns define a sector [28] that involves a group of coevolving residues critical for folding and enzymatic function, while local patterns involve isolated coevolving and contacting residue pairs outside the sector. Consistent with this distinction, the collective residues lie deeper in the protein structure (Fig.3A) and exhibit stronger functional effects upon mutation [25].

**Figure 3:**
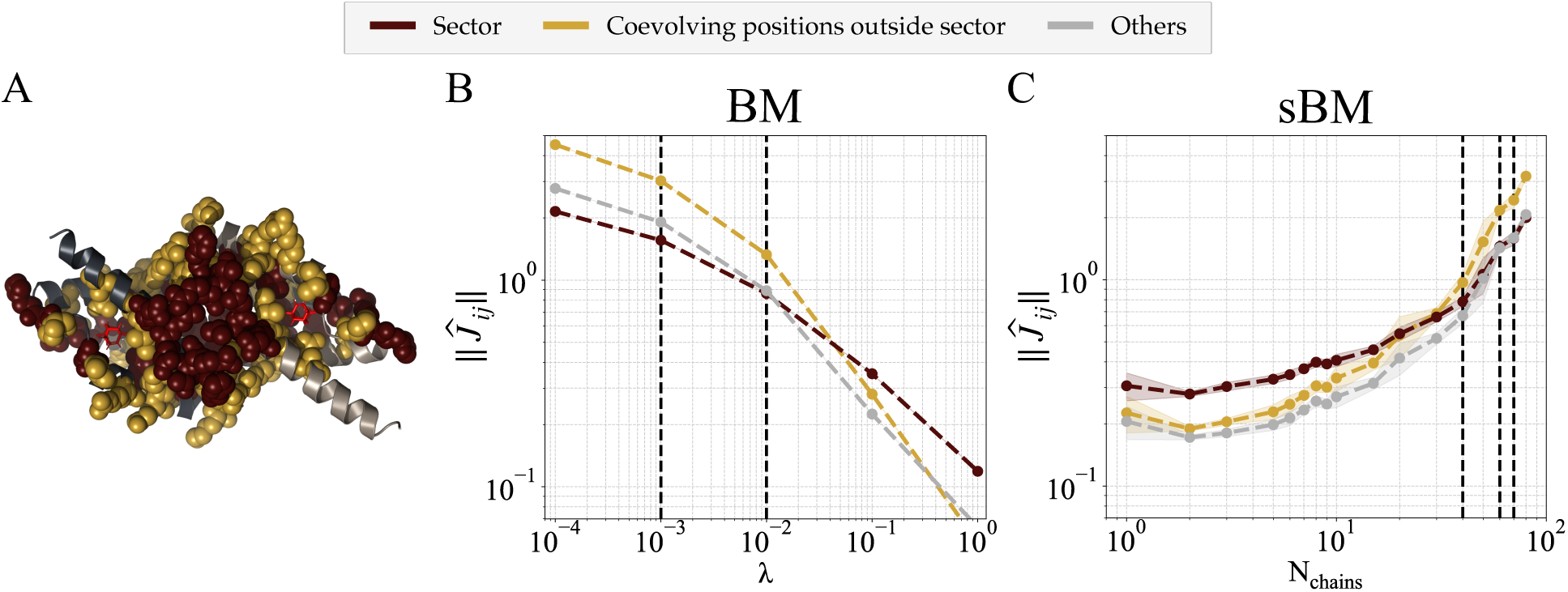
BM and sBM inference in the context of the chorismate mutase family. **A**. Representation of *E. coli* chorismate mutase structure, consisting of a dimer with two symmetric active sites (PDB: 1ECM). Sector positions represented by collective patterns of functional correlations and coevolving contacts outside the sector represented by isolated patterns of pairwise correlations (as defined by Kleeorin *et al.* [25]) are highlighted with yellow and red spheres, respectively. **B**. BM inference. Magnitude of inferred couplings as a function of *L*_2_ regularization (*λ* = *λ_h_* = *λ_J_* ). At low regularization, contact-associated couplings dominate; as regularization increases, this pattern reverses. Vertical dotted lines indicate the regularization levels used in Fig. 5. **C**. sBM inference. Magnitude of inferred couplings as a function of the hyperparameter *N*_chains_ (larger *N*_chains_ corresponding to weaker regularization). The relative importance of the two types of couplings as a function of *N*_chains_ mirrors the trend observed for the BM under increasing *L*_2_ regularization: couplings associated with local correlations dominate at large *N*_chains_, and this pattern reverses as regularization strengthens. However, consistent with the synthetic dataset, sBM regularization dampens coupling magnitudes far less sharply than *L*_2_ regularization. Vertical dotted lines mark the regularization level used in Fig. 5. In panels B and C, each point is the mean of 10 independent repeats. BM: average over 10 independently inferred BMs. sBM: average over 10 sBMs, where each sBM is itself the average of 10 independently inferred models. Error bars indicate the standard deviation across the repeats.

When inferred with BM, these patterns present biases that mirror those observed with synthetic data (Fig. 3B). The sBM shows a dependence of coupling magnitudes on *N*_chains_ similar to that of the BM on *L*_2_ regularization. Couplings associated with local patterns are stronger at high *N*_chains_, and this trend reverses as regularization increases. As for the synthetic dataset, the implicit regularization of the sBM affects coupling magnitudes less strongly than *L*_2_ regularization (Fig. 3C).

For real data, however, the true generative model is unknown, precluding a straightforward criterion for identifying an optimal regularization strength. As a consequence, the regularization yielding the highest fidelity in experiments cannot be inferred from Fig. 3C.

### Generativity in the context of synthetic data

According to Fig. 2, sBM enables the inference of models whose couplings more closely match those of the teacher model in an undersampled regime. We next examine whether this improved recovery of interactions translates into measurable gains in generative performance compared to models inferred using the BM. Generative performance must be evaluated along three complementary criteria: fidelity, novelty, and diversity [19]. For real protein data, fidelity is assessed experimentally by testing the functionality of generated sequences. In a mathematical setting, however, fidelity can be quantified directly from the teacher model by using the statistical energy of each sequence as a proxy for functionality. Here we consider a generated sequence to be functional if its statistical energy, computed using the teacher model, falls below a threshold determined from the energy distribution of sequences generated by the teacher itself (see Supplementary Methods and Fig. S9). Fidelity is then defined as the fraction of generated sequences classified as functional. Novelty is measured by computing, for each artificial sequence, its sequence distance to the closest training sequence (see Supplementary Methods and Fig. S10). Diversity is the fraction of training sequences that are identified as the closest training sequence from one of the generated sequences (see Supplementary Methods).

Figure 4 compares the generative performance of BM and sBM inferences across different regularization strengths. The goal is to obtain a model capable of generating artificial sequences with fidelity approaching that of the input model, while preserving levels of novelty and diversity comparable to those observed with the teacher model. In this synthetic setting, ∼90% of sequences generated with the teacher model are functional, the average distance to the nearest training sequence is ∼40%, and the diversity score is ∼58%.

**Figure 4:**
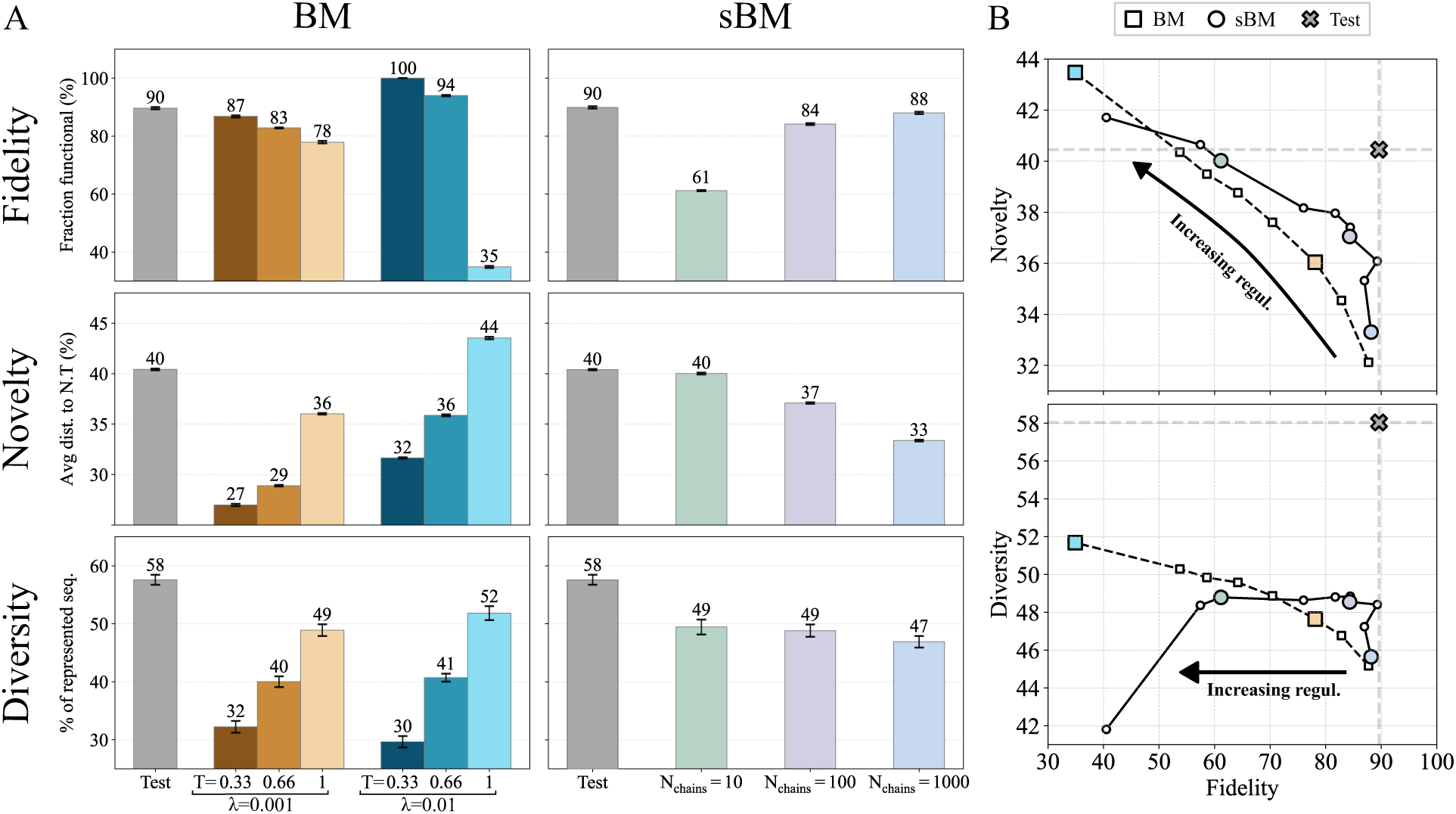
BM and sBM generative capacity in the context of synthetic data. **A.** Left panels show Fidelity, Novelty and Diversity scores for test sequences generated with the teacher model (grey) and sequences generated by two BM inferences inferred with *λ* = 0.001 (orange) and *λ* = 0.01 (blue), using sampling temperatures *T* ∈ {0.33, 0.66, 1}. Right panels show the same scores for test sequences (grey) and for sequences generated by three sBM inferences inferred with *N*_chains_ = 10 (green), 100 (purple), and 1000 (blue), all sampled at *T* = 1. **Fidelity:** Sequences are scored using the teacher model, and the fraction of functional sequences is defined as the proportion of sequences with a statistical energy below a threshold defined from the energy distribution of test sequences generated by the teacher model (see Supplementary Methods). **Novelty:** Novelty is quantified by computing, for each generated sequence, its sequence distance to the nearest training sequence. The novelty score corresponds to the average of this distribution (see Supplementary Methods). **Diversity:** Diversity is assessed using a coverage score defined as the fraction of training sequences that are the nearest neighbor of a generated sequence (see Supplementary Methods). **B.** Alternative representation of the same results, showing how there are trade-offs between the different criteria. Overall, sBM inference achieves a better compromise than BM inference (for clarity, only inferences at *T* = 1 are represented).

For BM with *L*_2_ penalties of 0.01 and 0.001, the fraction of functional sequences reaches 35% and 78%, respectively. Increasing fidelity toward the level observed for the teacher model can be achieved in two ways. One option is to further reduce regularization, which improves fidelity but inevitably leads to a loss of novelty (Fig. 4B). Alternatively, following Russ *et al.* [11], an additional hyperparameter can be introduced through the sampling temperature, such that sequences are generated according to 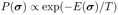. Sampling at *T <* 1 effectively rescales the model parameters, which lowers the statistical energies of generated sequences, and results in a substantially higher fraction of functional sequences. This improvement in fidelity, however, is associated with a marked reduction in both novelty and diversity, as shown in Fig. 4A.

sBM can achieve improved fidelity, novelty, and diversity without relying on low-temperature sampling. Figure 4A shows results obtained for three values of the regularization parameter, *N*_chains_ = {10, 100, 1000}. With weak regularization (*N*_chains_ = 1000), the inferred sBM generates a fraction of functional sequences close to that of the teacher model, with 88% of sequences classified as functional compared to 90% for the teacher model (the graph includes error bars reporting standard deviations over 10 independently generated synthetic datasets; these error-bar are barely visible because of their small magnitude, indicating that all reported differences are statistically significant). This gain in fidelity is accompanied by a reduction in novelty, with an average distance of 33% to the nearest training sequence, compared to 40% for the teacher model. In contrast, strong regularization (*N*_chains_ = 10) yields novelty levels comparable to those of the teacher model, but at the expense of reduced fidelity, with only 61% of generated sequences being functional on average. Consistent with the analysis of coupling-inference biases, the most favorable trade-off between fidelity and novelty is obtained for intermediate values of *N*_chains_. In particular, the model inferred with *N*_chains_ = 100 generates 84% functional sequences, with an average distance of 37% to the nearest training sequence. Importantly, this balance cannot be achieved using the BM under any choice of *L*_2_ regularization (see Fig. 4B). A comparable fidelity–novelty trade-off can be obtained by sampling at low temperature; however, this comes at the cost of a substantial loss of diversity (see Fig. 4A).

Altogether, these results demonstrate that sBM does not simply act by uniformly reducing parameter magnitudes or by rescaling energies. Moreover, the results obtained here with BM are representative of results obtained with other inference methods reported in the literature, notably arDCA [38] and adabmDCA [18] (Fig. S11 and Fig. S6). We also verify that reproducing the degree of correlation between the inferred and empirical statistics is not informative about generative performance, and even correlates negatively with novelty (Fig. S12).

### Generativity in the context of protein data

In the case of real data, assessing fidelity requires experiments to test the functionality of sequences generated by a model. In previous work on the chorismate mutase family [11], BM inferences were evaluated for two different values of the *L*_2_ regularization strengths, *λ* = 0.01 and *λ* = 0.001. In both cases, fewer than 5% of the generated sequences were functional. Introducing a sampling temperature *T <* 1, which rescales the model energy, substantially increased the fraction of functional sequences, reaching for example about 30% at *T* = 0.66 for *λ* = 0.01. As observed for synthetic data, this improvement in fidelity is obtained at the expense of reduced diversity and novelty (Fig. 5). In other words, the generated sequences become not only closer to natural proteins but also more similar to one another. This loss of diversity reflects a bias toward a restricted subset of sequences, which helps explain why the fraction of functional sequences at *T* = 0.33 can even exceed that observed in the natural dataset (Fig. S1).

**Figure 5:**
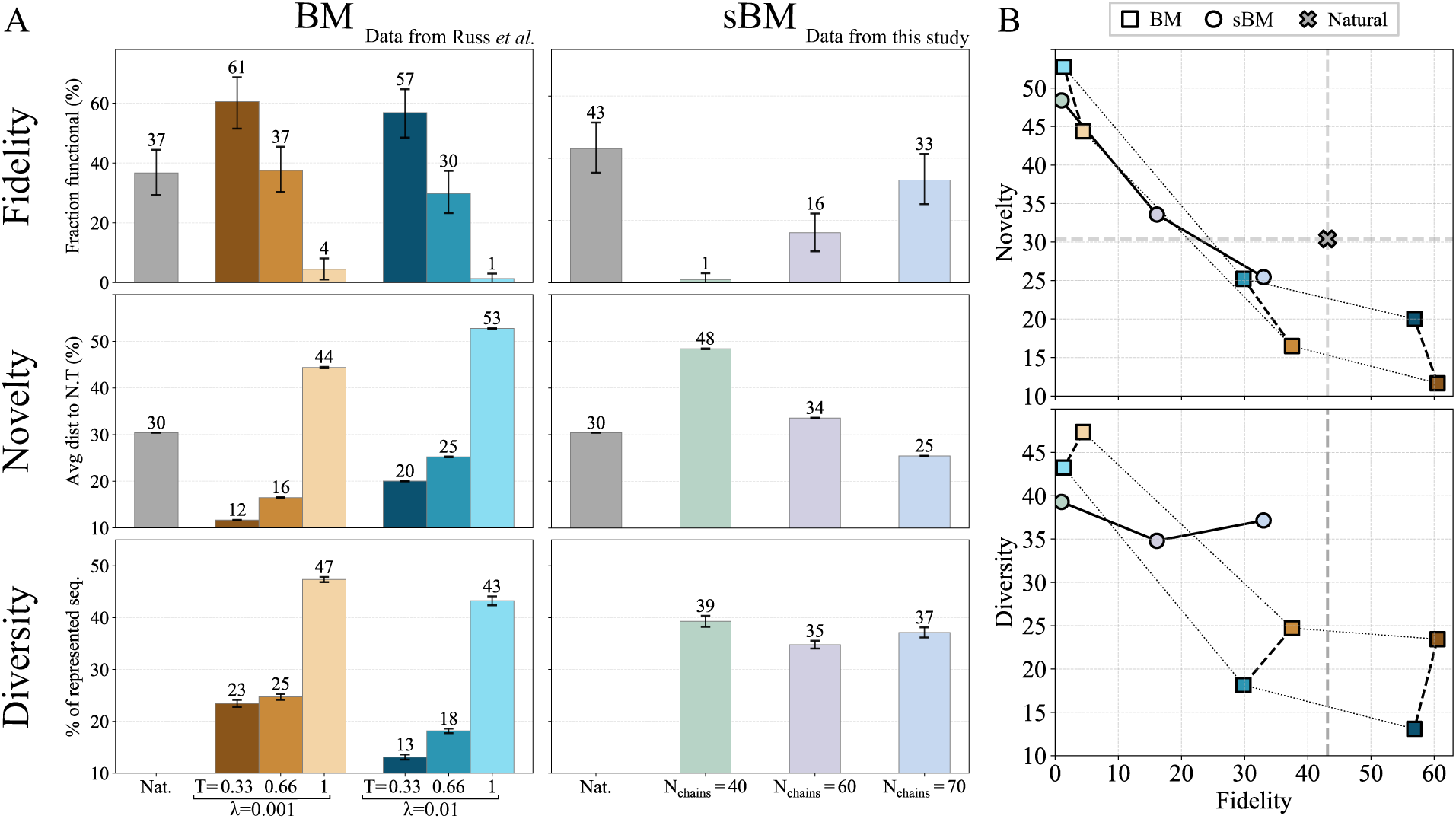
BM and sBM generative capacity in the context of the chorismate mutase family. **A.** Left panels show Fidelity, Novelty and Diversity scores for natural sequences (grey) and sequences generated by two BM inferences with *λ* = 0.001 (orange) and *λ* = 0.01 (blue), using sampling temperatures *T* ∈ {0.33, 0.66, 1}. Right panels show the same scores for natural sequences (grey) and for sequences generated by three sBM inferences with *N*_chains_ = 40 (green), 60 (purple), and 70 (blue), all sampled at *T* = 1. For each value of *N*_chains_, we independently inferred 50 sBM models (see Fig. S5) and averaged the inferred fields and couplings to obtain a single model. The experimentally characterized sequences were then generated from this averaged model. **Fidelity:** Artificial and natural sequences have been tested experimentally by Russ *et al.* [11] for the BM method and a similar protocol has been used in this study to test natural and artificial sequences generated with the sBM method (see Fig. S14 for comparison between natural CM relative enrichments in this study and Russ *et al.* data). **Novelty:** Novelty is quantified by computing, for each generated sequence, its sequence distance to the nearest training sequence. The novelty score corresponds to the average of this distribution (see Supplementary Methods). **Diversity:** Diversity is assessed using a coverage score defined as the fraction of natural sequences that are the nearest natural sequence of an artificial sequence (this metric is undefined for natural sequences). **B.** Alternative representation of the same results, showing how, overall, sBM can achieve a better trade-off between fidelity and novelty (top) and between fidelity and diversity (bottom). Here the results with BM at *T <* 1 are also represented.

To enable a direct comparison with the BM, we tested sBM inference with several regularization strengths corresponding to *N*_chains_ = {40, 60, 70}, three values chosen to achieve a novelty comparable to the original data (Fig. S4). Relative enrichment measurements were obtained for 1148 natural sequences, together with approximately 100 artificial sequences generated from the three sBM inferences (see Supplementary Methods and Fig. S13). The experimental results show that the fraction of functional sequences increases with *N*_chains_, that is, as regularization becomes weaker. This trend is consistent with the fact that reducing regularization leads to models that generate sequences which are, on average, closer to natural ones.

For *N*_chains_ = 70, the inferred sBM produces approximately 33% functional sequences, a level comparable to that obtained with BM inferences sampled at *T* = 0.66. However, the diversity score of the BM inferences at *T* = 0.66 falls below 25%, whereas the corresponding diversity score for the sBM inference is 37%. This result underscores the distinct nature of the sBM, which cannot be reduced to any BM with rescaled energies. The models inferred by the two approaches also differ by their distribution of statistical energies. For the BM inferences of Russ *et al.*, artificial sequences sampled at *T* = 1 exhibit, on average, higher statistical energies than natural sequences (Fig. S15), and lowering the sampling temperature shifts the energy distribution of artificial sequences downward. Although reducing the regularization strength can also yield artificial sequences with statistical energies comparable to those of the training set, these sequences display very limited novelty, remaining within 20% sequence distance of their closest natural neighbor (Fig. S16). In contrast, sequences generated by the sBM at *T* = 1 display statistical energies comparable to those of the training set (Fig. S15) while maintaining substantial sequence novelty, as desired.

We emphasize that different models may reproduce the empirical statistics equally well while having very different generative performance, as judged by their fidelity, novelty, and diversity (Fig. S17 and Fig. S18). We also emphasize that while applying a post hoc correction to rescale the energy function of BM models (low-temperature sampling) appears to improve fidelity in the context of chorismate mutases, this effect is confounded by experimental biases. The experimental assay indeed reports only certain regions of the sequence space as active, as revealed by its application to the natural sequences of the alignment [11]. Low-temperature sampling happens to overrepresent precisely these sections of the sequence space in experiments with the chorismate mutase family (Fig. S1). As a consequence, the fraction of active synthetic sequences obtained at low-temperature sampling is even larger than the fraction of active natural sequences. A different experimental assay or a different sampling of natural sequences can be expected to give different results. Irrespective of this, low-temperature sampling is necessarily at the expense of reduced novelty and diversity.

## Conclusion

Inferring generative models of proteins from alignments of homologous sequences exemplifies a pervasive challenge in biology: modeling high-dimensional systems of interacting elements from limited data. Proteins offer a particularly tractable case because their composition is precisely known, extensive sequence data are available, and functionality can be experimentally tested at high throughput. These advantages have enabled the design of novel, functional proteins whose sequences are far from any natural homolog [11]. Yet, like other generative models operating in high dimensions with limited data, protein models face an inherent trade-off between three competing goals [19]: fidelity (the fraction of functional proteins), novelty (their divergence from natural sequences), and diversity (the coverage of natural variation by designed sequences). A common way to address limitations from limited sampling is to introduce regularization. However, standard regularization schemes typically act uniformly on all model parameters, which can lead to major distortion effects: the resulting models may not only be approximate but systematically biased [25]. This issue becomes critical when the data contain statistical features over multiple scales, which is typical of biological systems. In proteins, these scales range from collective correlations defining functional sectors [28] to localized correlations reflecting structural contacts [3]. Uniform regularization distorts the relative representation of these two types of features.

Potts models provide a simple yet effective framework in which these issues are clearly apparent. Extensive studies in recent years have established common practices for assessing their accuracy, such as reproducing statistics, and for addressing undersampling through early stopping, L2 regularization, and pseudocounts. However, few studies have tested the generative capabilities of these models. In a notable exception, the model was found to generate mostly nonfunctional sequences [11]. Applying a post hoc uniform rescaling of parameters provided a pragmatic workaround, increasing the fraction of functional generated sequences [11]. Yet, this approach is not a panacea. First, it reduces the novelty and diversity of generated sequences. Second, the apparent improvement in fidelity in the tested case arises, at least in part, because the reduced diversity coincides with a bias in the experimental assay (Fig. S1). Our work demonstrates that this reduction in diversity is not inevitable: a model can be inferred that, when sampled without rescaling, achieves comparable fidelity while preserving diversity to a greater extent.

To achieve these results, we introduced an inference method, termed sBM, that employs an alternative regularization strategy to more accurately represent features across scales. This strategy relies on three key ingredients, each of which we find to be necessary: an approximate second-order optimization method (the L-BFGS algorithm with a small value of the parameter *m* controlling its approximation), early stopping (a limited number *N*_iter_ of iterations), and limited sampling of the model statistics (a limited number *N*_chains_ of Monte Carlo samples). While the latter two ingredients are common to other implementations of BM, which are also constrained by finite values of *N*_chains_ and *N*_iter_, our criteria for selecting these values differ and lead us to distinctly different choices. Rather than focusing on reproducing the statistics of the data, a poor proxy for generativity, we prioritize three criteria widely recognized as key to generative models: fidelity (the fraction of functional generated sequences), novelty (the difference between generated and natural sequences), and diversity (the coverage of natural variation by generated sequences). With limited data, these three properties involve trade-offs. In the context of proteins, novelty and diversity can be estimated computationally, but fidelity requires experimental validation. In practice, we set *m* = 1, determine *N*_iter_ based on a plateau in the iteration dynamics, and constrain the values of *N*_chains_ to be tested experimentally based on novelty. We demonstrated the effectiveness of our approach using both synthetic data with known ground truth and experimental tests on chorismate mutases. We also verified that alternative methods, such as first-order gradient descent or persistent contrastive divergence, other regularization parameters (e.g., pseudocounts), or alternative criteria for setting the same regularization parameters we use, fail to capture the multiscale statistics of the data as well and to achieve a comparable trade-off between fidelity, novelty and diversity.

Like many machine learning methods, our approach is guided by heuristic rather than formal arguments, and its value ultimately rests on empirical validation. Nevertheless, our results provide both a proof of principle that further progress is possible and a direction for achieving it: generative models can be improved by striving for a more balanced representation of statistical features across scales. Because high-dimensional, multiscale systems inferred from limited data are ubiquitous in biology, these insights should extend beyond proteins to a broad range of biological modeling problems.

## Acknowledgments

EH acknowledges funding from NSF Graduate Research Fellowship (NSF 2140001). OR acknowledges funding from ANR-21-CE45-0033.

## Resource availability

### Lead Contact

Information and requests for resources and reagents should be directed to the lead contact

### Data and code availability

- The experimental data will be made available upon publication.
- Codes for BM and sBM inferences are publicly available at https://github.com/StatBio/Stochastic-Boltzmann-Machine.git.

## Supplementary Methods

### Teacher model and synthetic data

The teacher model is identical to the one introduced by Kleeorin *et al.* [25]. In particular, the input coupling matrix *J* ^inp^ is designed to encode isolated patterns, a medium-size collective pattern, and a large collective pattern. The model, illustrated in Fig. 1A, consists of *L* = 20 positions with *q* = 10 possible amino acids at each site. It includes zero fields, and all non-zero couplings are set to the same interaction strength 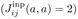. This choice favors identical amino acids at coupled positions and eliminates frustration.

Training data are generated by sampling *M* = 300 synthetic sequences from the teacher model using Markov chain Monte Carlo with the Metropolis–Hastings algorithm. Samples are obtained after 10^5^ MCMC steps, starting from independent random initial sequences.

### Multiple sequence alignment of the chorismate mutase family

The alignment of the chorismate mutase family is as described by Kleeorin *et al.* [25]. It contains 1258 sequences and 96 positions. In [11], the authors do not set apart a test dataset. We similarly use all sequences for sBM inferences.

### Collective vs. isolated patterns in the chorismate mutase family

The sector, represented by collective patterns of functional correlations, and the coevolving contacts outside the sector, represented by isolated patterns, considered in this work were previously defined by Kleeorin *et al.* [25] as follows:

- The sector is defined as top-ranked pairs inferred with strong regularization in the absence of average product correction (APC) that belong to the functional network of mutationally sensitive positions, while not corresponding to structural contacts.
- The coevolving contacts outside sector are identified as top-ranked pairs inferred in the low-regularization limit with APC that correspond to contacts in the three-dimensional structure, but do not belong to the functional network of mutationally sensitive positions.

### Model inference

For both BM and sBM inferences, parameters are estimated by minimizing the average negative log-likelihood using gradient-descent optimization:

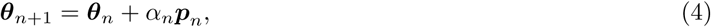

where ***p****_n_* denotes the search direction, *α_n_*the learning rate, and ***θ****_n_* is the parameter vector at iteration *n*. As detailed in Algorithm 1, the computation of the search direction ***p****_n_* depends on the inference method (see the two following sections for further details on each inference procedure). However, the following implementation choices are common to both BM and sBM inferences:

- Empirical statistics are weighted to mitigate phylogenetic bias [43]: the weight of a sequence is inversely proportional to the number of sequences sharing more than *r*% identity with it; we set *r* = 70%.
- No pseudocount is added to the empirical statistics.
- All couplings and fields are initialized to zero.
- Model statistics are approximated by Monte Carlo sampling using *N*_chains_ sequences. For all results presented in this paper, simulations are initialized randomly, and the chains are generated with *k* = 10^5^ steps.

#### Algorithm 1: Gradient-based optimization

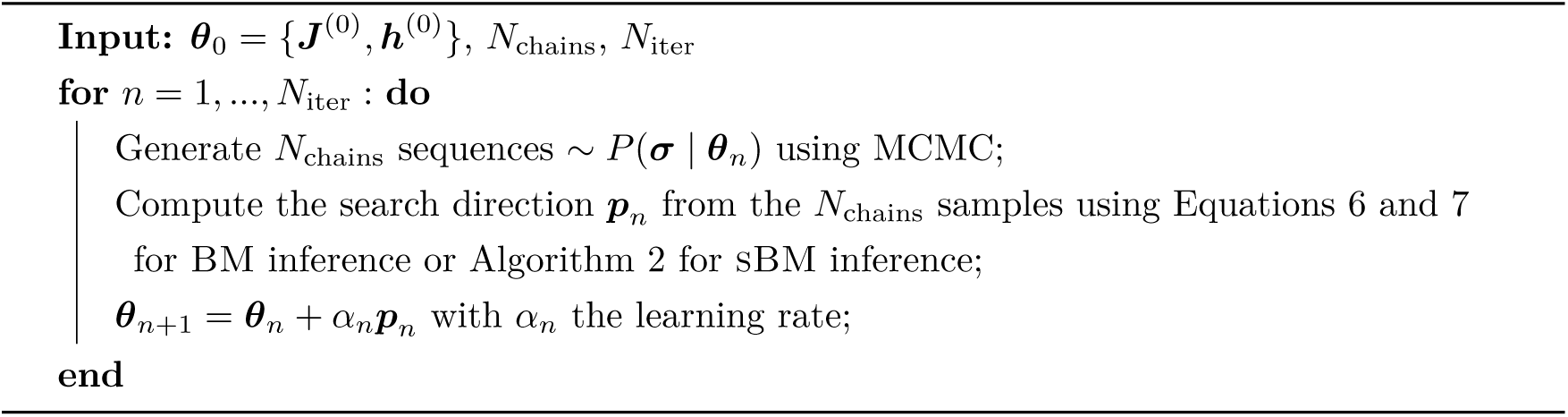

### BM inference

The BM inference is performed using vanilla gradient descent, such that the search direction ***p****_n_* is determined by the gradient of the objective function *g*(***θ****_n_*):

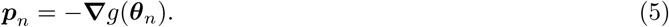

The gradient of the negative log-likelihood with *L*_2_ regularization is given, respectively, for the fields and the couplings by

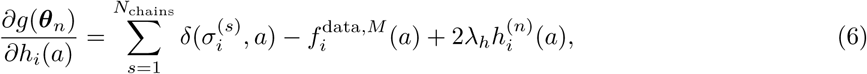

and

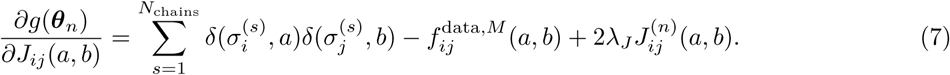

Unless otherwise stated, the number of Monte Carlo samples *N*_chains_ used to approximate the model statistics at each step of the gradient descent is set to 1000. Finally, the parameters are updated according to Eq. 4 using a decreasing learning rate *α_n_* = (*n* + 1)*^−^*^0.2^.

### sBM inference

The sBM inference is done using Limited-memory BFGS [37] which is a variant of BFGS developed by Broyden, Fletcher, Goldfarb, and Shannon. Both are quasi-Newton optimization algorithms taking the form

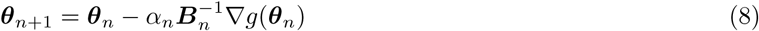

with ***θ****_n_* the current parameter estimate, *α_n_* the learning rate, *g* the objective function (negative log-likelihood) and ***H****_n_* = ***B_n_****^−^*^1^ an approximation of the inverse Hessian. BFGS incrementally updates an estimate of the inverse Hessian which requires 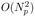 space and time per iteration, *N_p_* being the number of parameters. In L-BFGS, the inverse Hessian is approximated using only the last *m* optimization steps, which is particularly advantageous when the storage cost of the Hessian is excessive. In our implementation, we use a simple and efficient procedure to compute the search direction ***p****_n_* = −***H****_n_***∇***g*(***θ****_n_*), based on the *m* most recent pairs of vectors {***s****_n_* = ***θ****_n_*_+1_ − ***θ****_n_,* ***y****_n_* = **∇***g*(***θ****_n_*_+1_) − **∇***g*(***θ****_n_*)} stored during the optimization [44] (see Algorithm 2).

In general, learning rates must be chosen using a line search procedure to ensure that parameter updates satisfy the Wolfe conditions. However, this requires multiple evaluations of the objective function, resulting in a computational cost that is prohibitive in our context. In our implementation, the initial learning rate is set to *α*_0_ = ∥**∇***g*(***θ***_0_)∥*^−^*^1^, and the first search direction is taken as ***p***_0_ = −**∇***g*(***θ***_0_). For all subsequent iterations, the learning rate *α_n_*is fixed to 1.

If, at any iteration, the condition 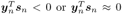 is met, the inverse Hessian estimate is not updated and *H_n_*_+1_ = *H_n_*.

#### Algorithm 2: Calculation of ***p****_n_* = −***H****_n_***∇***g*(***θ****_n_*)

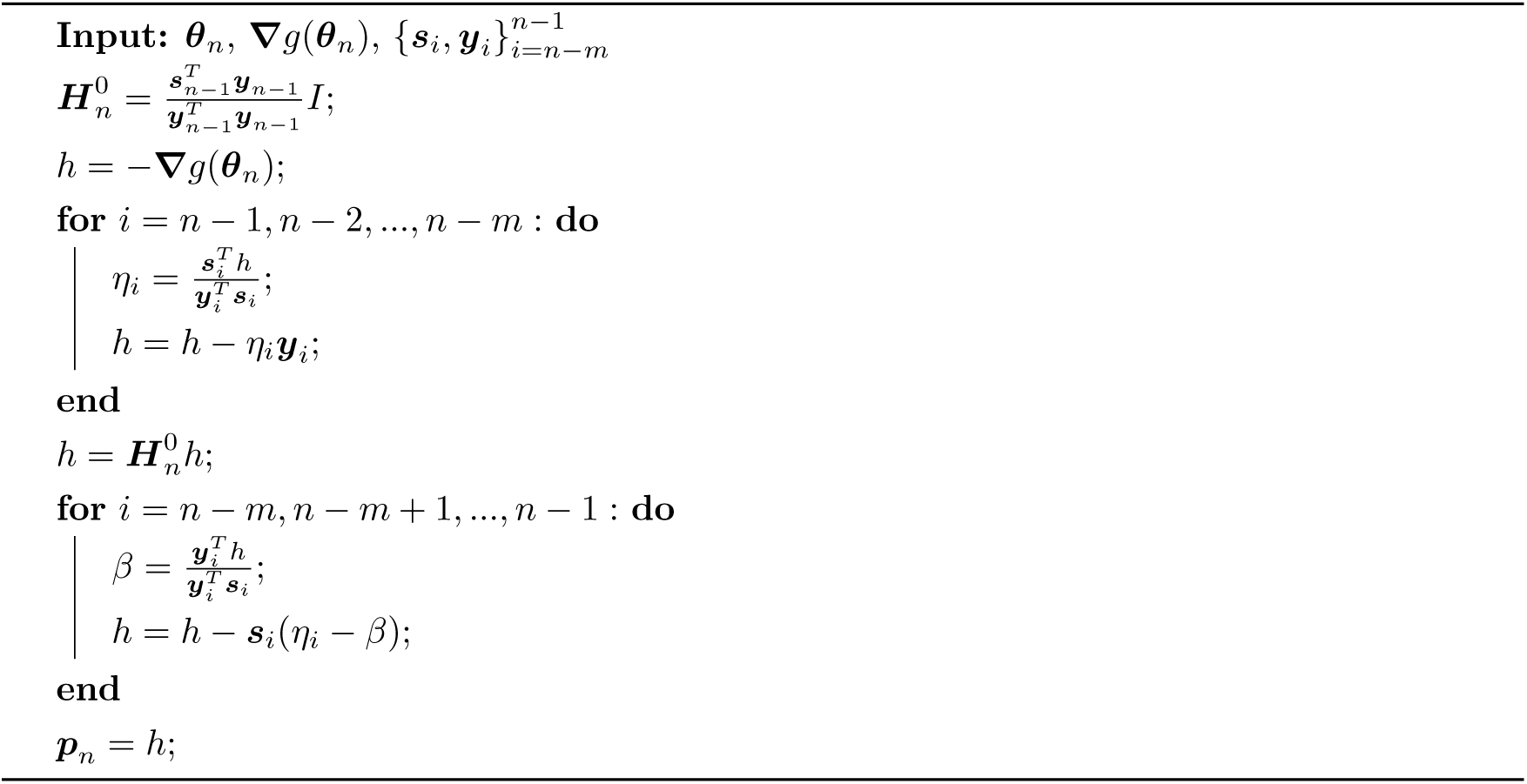

Selecting hyperparameters associated with regularization is non-trivial, particularly given the unsupervised nature of the task. In practice, for the chorismate mutase dataset we set *m* = 1. To determine the number of iterations, we monitor the average Frobenius norm of the inferred couplings throughout the optimization and choose *N*_iter_ such that the procedure is stopped once this quantity has reached a plateau (here *N*_iter_ ∼ 400). Finally, the parameter *N*_chains_ controls the regularization strength. As illustrated in Supp. Fig. S5, varying *N*_chains_ changes the plateau value reached on average by the inference, and directly impacts the novelty of the generated sequences (Fig. 5). The inference also exhibits variability across runs despite fixed hyperparameters. To account for this variability, each reported sBM model was obtained by averaging the inferred fields and couplings over 10 independent inference runs. For the three models selected for experimental characterization, the average was computed over 50 independent inference runs.

### Gauge invariance

Because the number of parameters exceeds the number of independent statistical constraints, the model is overparameterized. This overparameterization gives rise to a gauge invariance, meaning that the probability distribution remains unchanged under a certain class of transformations, referred to as gauge transformations. In the presence of L2 regularization, this gauge degeneracy is broken during inference, since the L2 penalty favors a particular parameterization among gauge-equivalent representations of the same model.

For all plots involving inferred parameters, results are given in the zero-sum gauge, obtained as

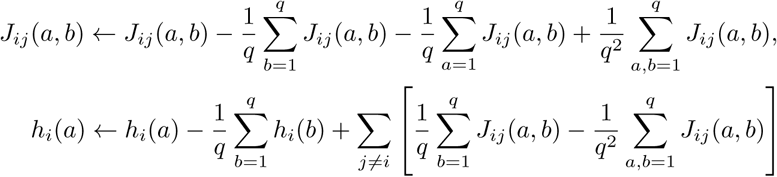

This choice of gauge minimizes the Frobenius norm of the coupling matrix by imposing

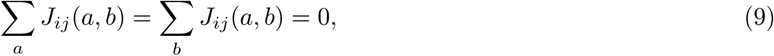

while the fields satisfy

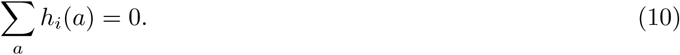

### Sampling of artificial sequences

Artificial sequences are obtained by sampling with the same MCMC algorithm used for the inference. The number of MCMC steps *k* is chosen to be the same as the one used during the inference, i.e. 10^5^ in this study.

### Fidelity score in the context of synthetic data from a mathematical model

In the teacher-student setting, the fidelity of generated sequences is quantified directly using the teacher model. To this end, synthetic datasets are generated from both the teacher model and the inferred models, and all sequences are evaluated using the energy function of the teacher model.

Specifically, 10,000 sequences are sampled from the teacher model (referred to as test sequences) and 10,000 sequences are sampled from the inferred model. Sequence sampling is performed using the same MCMC procedure as described above, with identical sampling parameters across models. For each sequence, the statistical energy is computed using the parameters of the teacher model. Sequences are then classified as functional based on their statistical energy. A functionality threshold is defined from the energy distribution of test sequences generated by the teacher model, as shown in Fig. S9. Sequences with energies below this threshold are considered functional, while sequences with higher energies are classified as non-functional.

The fidelity of a model is quantified as the fraction of generated sequences classified as functional. To estimate variability due to finite sampling, the entire procedure is repeated independently ten times for each model and each set of hyperparameters. Reported fidelity values correspond to averages over these repeats, and error bars represent the associated standard deviations.

### Fidelity score in the context of protein data

In the context of protein data, the functionality of each generated sequence is estimated from an experimental assay, as described below.

To estimate the uncertainty associated with the finite number of experimentally characterized sequences, fidelity is computed using bootstrap resampling. Specifically, 100 sequences are sampled with replacement from each dataset, and the fraction of functional sequences is computed for each bootstrap sample. This procedure is repeated 5,000 times. The reported fidelity corresponds to the mean of the resulting bootstrap estimates, while the error bars represent their standard deviation.

### Novelty score

Novelty quantifies how distinct the generated sequences are from the training data. Artificial sequences are generated from the inferred models, and for each artificial sequence its distance to the closest sequence in the training set is computed. For two sequences ***σ*** and ***σ****^′^* of length *L*, the Hamming distance is defined as

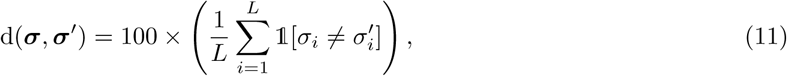

where 1[·] denotes the indicator function. In this calculation, we choose not to count positions where both sequences contain a gap (but including gap-gap matches does not qualitatively affect the results).

For each artificial sequence, the distance to its nearest training sequence is computed, and the novelty score is defined as the average of these distances over a set of 10,000 generated sequences. This procedure is repeated independently ten times, and the reported novelty score corresponds to the mean across repeats, with error bars given by the standard deviation.

In the teacher-student setting, the same procedure is applied to sequences generated from the teacher model, providing a reference level of novelty expected in the absence of inference errors. For real protein data, the novelty score reported for natural sequences is computed by considering, for each natural sequence, its distance to the closest other natural sequence in the dataset, excluding self-comparisons.

### Diversity score

The goal of the diversity metric is to assess how well artificial sequences cover the variability present in the training dataset.

For each inferred model, a number of artificial sequences equal to the size of the training dataset is generated. For each artificial sequence, the closest natural sequence in the training set is identified based on sequence distance. A natural sequence is then considered *represented* if it is identified as the nearest natural neighbor of at least one artificial sequence. The diversity score is defined as the fraction of natural sequences that are represented in this way. This procedure is repeated independently ten times for each model. The reported diversity score corresponds to the mean over these repetitions, and error bars represent the associated standard deviation.

In the teacher–student setting, this diversity metric is also computed for sequences generated directly from the teacher model. In this case, artificial sequences are generated from the teacher model and compared to training sequences sampled from the same distribution. This provides a reference value for the expected diversity in the absence of inference errors and serves as a baseline for comparison with inferred models.

### Chorismate mutase library construction

Genes encoding the artificial protein sequences were ordered as a mixed pool of single-stranded oligonucleotides (IDT oPools). A nucleotide sequence corresponding to each designed amino acid sequence was determined by randomly sampling codons proportionally to their usage in K12 *Escherichia coli* [45]. Two stop codons (TAATGA) were added to the end of the coding sequence to minimize read-through. The coding sequence and stop codons are flanked by universal primer and restriction sites: a primer site for GSP2 (5’-AAACCGGAGCCATACAGTAC-3’) and an NdeI restriction site (5’-CATATG-3’) precede the 5’ end of the coding sequence, and the stop codons at the 3’ end of the coding sequence are followed by an XhoI restriction site (5’-CTCGAG-3’) and a reverse complement priming site for GSP107 (5’-CCGTGCGACAAGATTTCAAG-3’) [46]. No internal priming sites for GSP2 or GSP107 (defined as a 10 bp or more match on either strand), internal restriction sites, or internal repeated subsequences of 10 bp or more were present in the final oligonucleotides. No more than 9 repeated A or repeated T bases and no more than 5 repeated G or repeated C bases were allowed. The GC content of both the coding sequence and the full oligonucleotide was restricted to be between 40% and 60%. The GC content of the first 60 (5’) bases was restricted to be less than 65%. All 100 bp windows of the oligonucleotide had a GC content below 70%, all 20 bp windows had a GC content below 90%, and the highest and lowest GC content within any 50 bp window differed by no more than 50%.

Oligonucleotide pools were amplified by PCR using KAPA HiFi DNA polymerase (Roche) with 1X KAPA HiFi Fidelity buffer, 0.2 mM dNTPs, 2 mM GSP2, 2 mM GSP107, 0.02 *µ*L KAPA HiFi DNA polymerase, and 0.4 ng/*µ*L template DNA in a final reaction volume of 50 *µ*L. The reaction was incubated at 95*^◦^*C for 3 min (polymerase activation), followed by 14 cycles of 98*^◦^*C for 20 s (denaturation), 60*^◦^*C for 15 s (annealing), and 72*^◦^*C for 15s (extension). After 14 cycles, a final extension was performed by incubating the reaction at 72*^◦^*C for 2 min. To separate the desired PCR products from DNA synthesis artifacts, amplicons of the target size (300-350 bp) were gel purified and concentrated (Zymo Research). Purified amplicons were digested with NdeI and XhoI (NEB), ligated into NdeI- and XhoI-digested pKTCTET-0 [47], column purified (Zymo Research), and transformed into XL1-Blue *E. coli* (Agilent) to yield *>*1000X coverage per gene. The entire transformation was cultured in 100 mL LB media containing 100 *µ*g/mL sodium ampicillin at 37*^◦^*C overnight, after which plasmids were purified via miniprep (Macherey Nagel).

### Chorismate mutase selection assay

A previously reported protocol was used to carry out the selection assay [11, 47]. The library of sBM designs in pKTCTET-0, a previously constructed library of natural CM coding sequences in pKTCTET-0 [11], and a previously constructed standard curve library of *E. coli* CM point mutants in pKTCTET-0 [11] were each diluted to a concentration of 0.1 ng/mL and separately transformed into the CM-deficient assay strain KA12/pKIMP-UAUC [48] to yield *>*1000X coverage per gene. Transformation mixtures were incubated for one hour at 37*^◦^*C immediately following transformation, then diluted separately into 500 mL LB media containing 100 *µ*g/mL sodium ampicillin and 30 *µ*g/mL chloramphenicol and incubated overnight at 30*^◦^*C. Aliquots of each overnight culture maintaining *>*1000X coverage of each gene were supplemented with 20% glycerol and frozen at -80*^◦^*C.

Glycerol stocks of KA12/pKIMP-UAUC carrying each library were revived separately in LB media containing 100 *µ*g/mL sodium ampicillin and 30 *µ*g/mL chloramphenicol overnight at 30*^◦^*C. Overnight cultures were used to inoculate separate cultures of M9c minimal media [47] supplemented with 100 *µ*g/mL sodium ampicillin, 30 *µ*g/mL chloramphenicol, 20 *µ*g/mL of L-phenylalanine (F), and 20 *µ*g/mL of L-tyrosine (Y) (M9cFY, non-selective conditions) to an OD_600_ of 0.045. M9cFY cultures were grown to 30*^◦^*C to an OD_600_ of approximately 0.2. Cultures were mixed to yield equal representations of each gene in the designed and natural libraries, and 4X representation of each gene in the standard curve. The mixed culture was washed twice with M9c media (no FY). The washed culture was used to inoculate 500 mL M9c (selection) and 500 mL M9cFY (non-selection control), each supplemented with 100 *µ*g/mL sodium ampicillin, 30 *µ*g/mL chloramphenicol, and 3 ng/mL doxycycline hyclate (to induce gene expression) to an OD_600_ of 0.001. The remainder of the mixed culture was harvested and resuspended in 2 mL LB with 100 *µ*g/mL sodium ampicillin and 30 *µ*g/mL chloramphenicol, grown overnight at 37*^◦^*C, and harvested for plasmid purification (the input sample) via miniprep (Macherey Nagel). The selection and non-selection cultures were grown for 24h at 30*^◦^*C, passaging as necessary to maintain an OD_600_ *<* 0.1. For both the selection and non-selection cultures, 50 mL of culture was harvested, resuspended in 2 mL LB with 100 *µ*g/mL sodium ampicillin and 30 *µ*g/mL chloramphenicol, grown overnight at 37*^◦^*C, and harvested for plasmid purification via miniprep.

Input, selected, and non-selected samples were amplified using two rounds of PCR with Q5 High-Fidelity DNA polymerase (NEB) to add adapters and indices for Illumina sequencing. The first round of PCR (2 cycles) added 6-9 random nucleotides (for initial focusing during sequencing) and part of the i5 and i7 Illumina adapters. The products of the first round of PCR were purified via bead-cleanup (Beckman Coulter). The second round (18 cycles) added the remaining i5 and i7 adapters as well as TruSeq indices to each sample. In addition to a low number of total cycles, a high initial DNA concentration in both rounds was used to minimize amplification bias. Second-round PCR products were gel purified (Zymo Research), quantified via Qubit (ThermoFisher), mixed in equimolar ratios, diluted to 4 nM, and sequenced on an Illumina MiSeq system with a 2×300 paired-end kit.

### Experimental data processing

Paired-end Illumina reads were joined using FLASH [49], trimmed to the NdeI and XhoI restriction sites, and translated. For the designed and natural libraries, only exact matches to library variants were counted. For the standard curve point mutants, only exact match sequences with a correct 4 bp barcode were counted. For natural and designed variants, sequences with fewer than 11 reads in the input sample were excluded from analysis. After filtering for low initial count, a pseudocount of 1 read per sequence per sample was added for calculations. Relative enrichment scores (r.e.) for the selected sample were calculated as

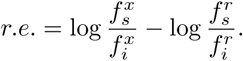

The variables 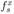 and 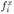 are the frequencies of each allele *x* in the selected (*s*) or input (*i*) samples. The variables 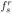 and 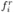 are the corresponding values for the *E. coli* chorismate mutase, the reference allele. For comparison to published data, normalized relative enrichment scores were calculated such that a value of 1 corresponds to the *E. coli* wild-type and a value of 0 corresponds to the mean of the near-null mode across all libraries as fit by a two-mode Gaussian mixture model.

Functional labels to evaluate fidelity were assigned according to a two-mode Gaussian mixture model fit only to the natural CM relative enrichment scores, as in previously published experiments [11]. If the mixture model predicted a relative enrichment score as more likely drawn from the high relative enrichment mode, the corresponding sequence was assigned a positive (functional) label.

## Supplementary figures

**Figure S1:**
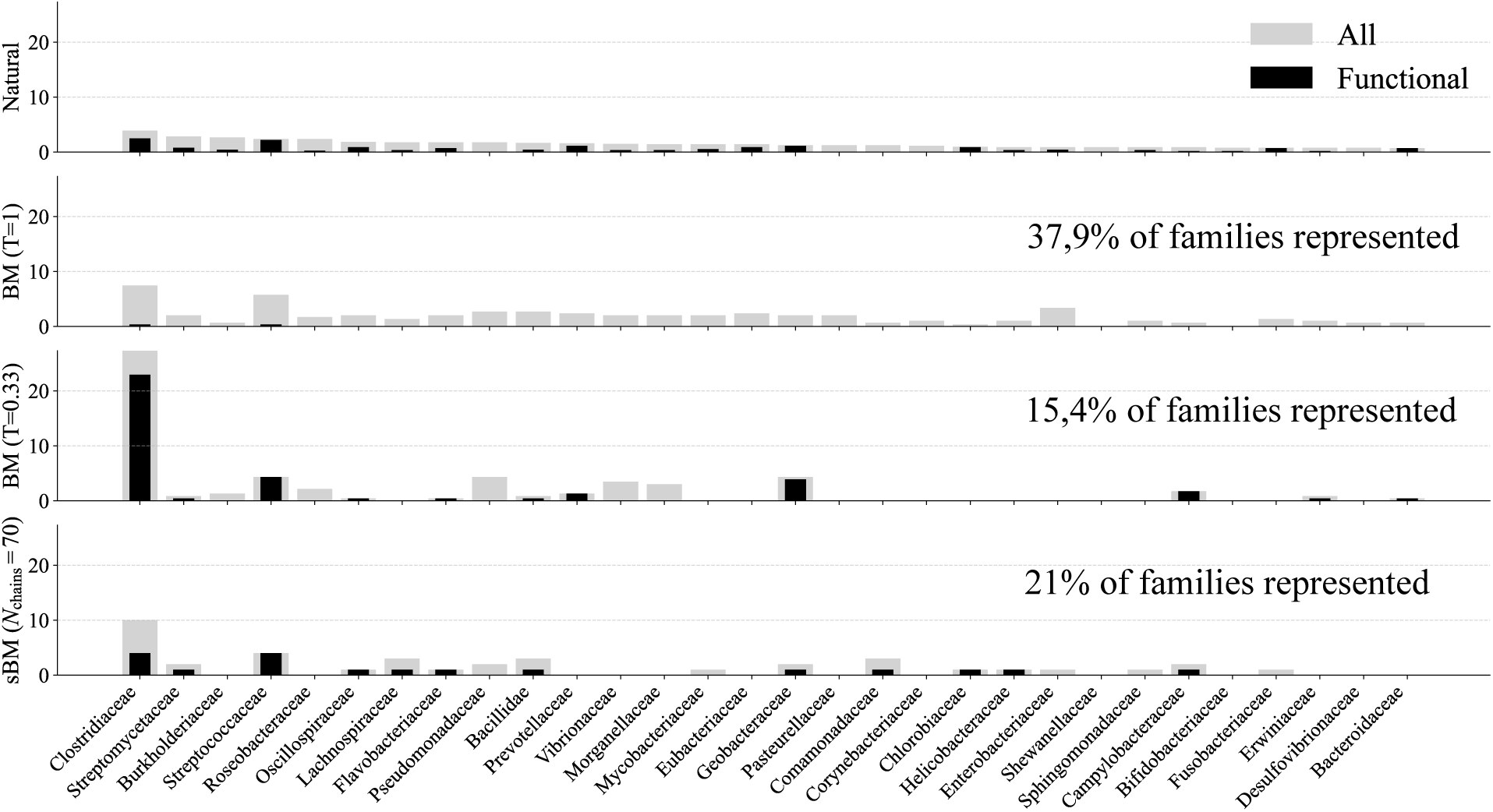
CM family: Effect of temperature rescaling on the phylogenetic distribution of generated sequences. The top panel shows the distribution of natural sequences among the 20 most represented phylogenetic families in the chorismate mutase MSA. Gray bars indicate the proportion (%) of sequences belonging to each family, while black bars show the proportion of sequences that the experimental assay reports as active. The lower panels display the corresponding distributions for sequences generated by the BM model of Russ *et al.* [11], inferred with an L2 regularization strength of *λ* = 0.01, and sampled at *T* = 1 (middle) or *T* = 0.33 (bottom). Each generated sequence was assigned to the family of its closest natural neighbor. Lowering the sampling temperature biases generation toward a subset of phylogenetic families, explaining the increased fraction of functional sequences observed at *T* = 0.33 as arising from an alignment between the low-temperature sampling bias and the experimental bias.

**Table 1:** Summary of sequences complementing growth in CM-null *E. coli* with a minimum count of 11 in the input library. Results are shown for the natural sequence library and for sequences generated by sBM models inferred with *N*_chains_ = 40, 60, and 70. Sequences are classified as functional or non-functional using either a threshold defined as +3*σ* relative to the mean of a null mode fitted on the full dataset, or a two-mode Gaussian mixture model trained on natural sequences only.

| Library | # seqs | # functional<br>($+3\sigma$ threshold) | # non-functional<br>($+3\sigma$ threshold) | # functional<br>(2-mode GMM<br>trained on Nats) | # non-functional<br>(2-mode GMM<br>trained on Nats) |
| --- | --- | --- | --- | --- | --- |
| Natural Library | 1148 | 531 (46%) | 617 (54%) | 493 (43%) | 655 (57%) |
| sBM $N_{\text{chains}}=40$ | 99 | 1 (1%) | 98 (99%) | 1 (1%) | 98 (99%) |
| sBM $N_{\text{chains}}=60$ | 99 | 19 (19%) | 80 (81%) | 16 (16%) | 83 (84%) |
| sBM $N_{\text{chains}}=70$ | 100 | 36 (36%) | 64 (64%) | 33 (33%) | 67 (67%) |

**Figure S2:**
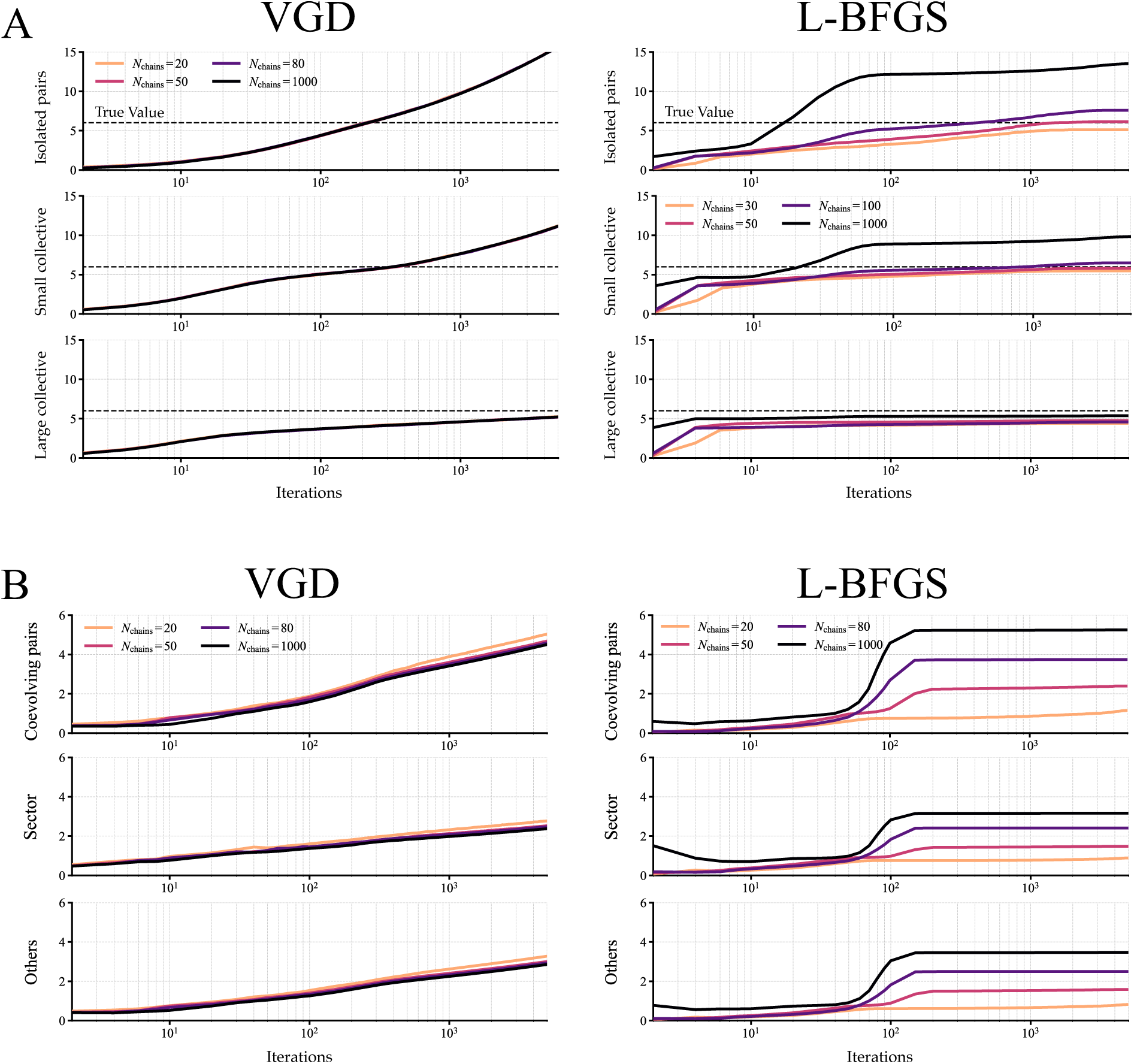
Interaction between implicit regularization and the optimization algorithm. Evolution of the average inferred couplings as a function of the optimization iterations. In both panels, the left and right columns correspond to Vanilla Gradient Descent (VGD) and L-BFGS optimization, respectively. Each panel consists of three rows showing the average inferred coupling strengths for different categories of couplings, while the different colors correspond to different values of *N*_chains_. **A.** Synthetic data. The three rows correspond to isolated pairs, small collective units, and large collective units. Dashed horizontal lines indicate the corresponding ground-truth coupling strengths. Under VGD, varying *N*_chains_ from 10 to 1000 has essentially no effect on the inference dynamics, whereas under L-BFGS it strongly affects the convergence and the inferred parameters. **B.** Protein data. The three rows correspond to coevolving pairs outside sectors, within sectors, and all remaining pairs (as defined by Kleeorin *et al.* [25]). As in the synthetic setting, L-BFGS makes the inference dynamics strongly dependent on *N*_chains_, whereas VGD remains weakly sensitive to this hyperparameter.

**Figure S3:**
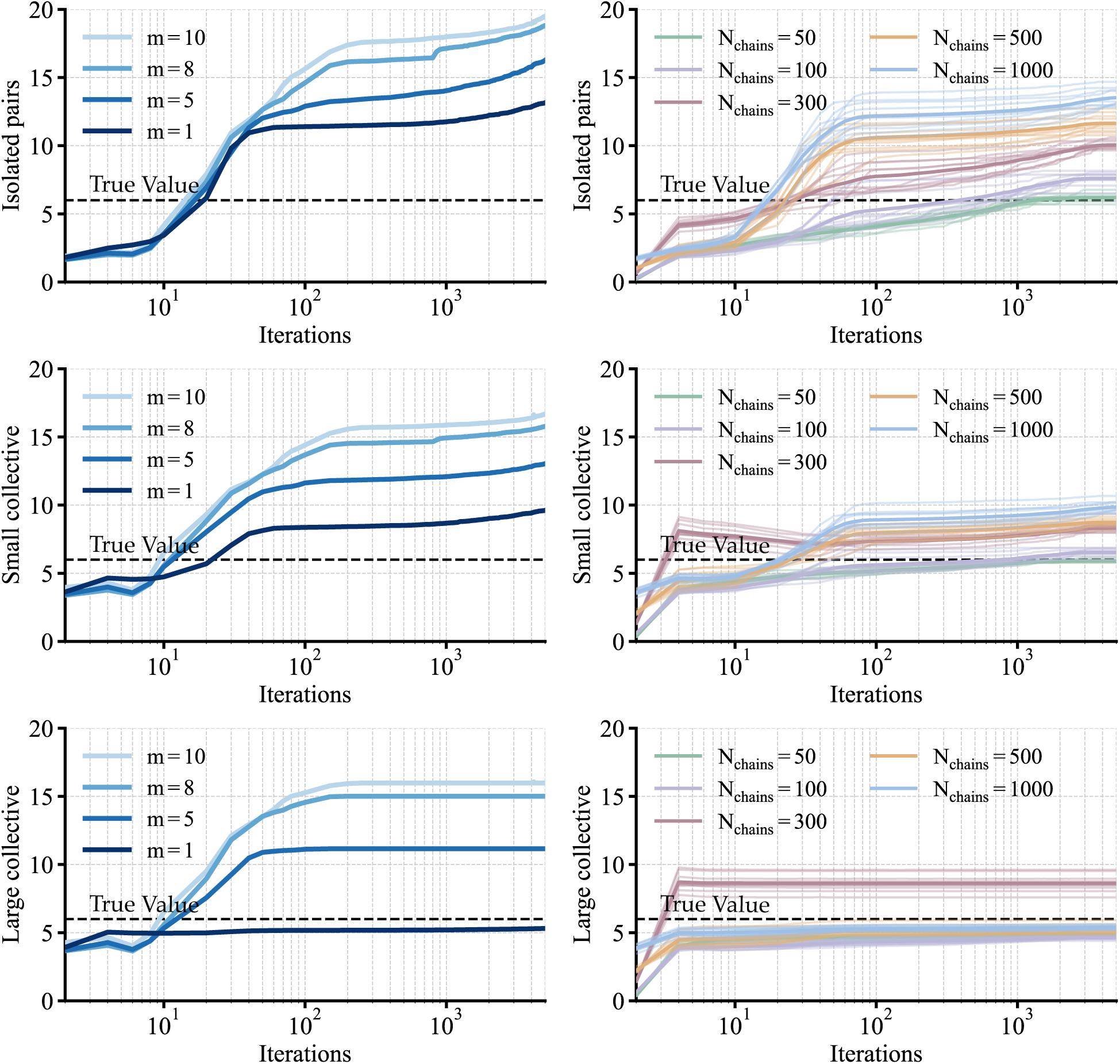
Synthetic data: sBM learning as a function of the hyperparameters *m* **and** *N*_chains_. Each panel shows the evolution of the average coupling magnitude as a function of the number of optimization iterations, for different types of couplings in the teacher model. The dashed black line indicates the true value of the teacher model. **Rows:** From top to bottom: isolated pairwise couplings, small collective couplings, and large collective couplings. **Left column:** Effect of the L-BFGS memory parameter *m*, with *N*_chains_ = 1000 fixed. The learning dynamics for the couplings appear to exhibit several distinct phases. The first phase, spanning roughly the initial 10 iterations, shows minimal progress. This is followed by a second phase of accelerated convergence, which is notably influenced by the choice of *m*: smaller values of *m* lead to an earlier termination of this acceleration phase. The learning dynamics then enter a plateau lasting until approximately iteration 1000, after which the inferred couplings begin to increase again. While only the first 5000 iterations are shown, the couplings continue to grow beyond that point due to the absence of regularization. **Right column:** Effect of the number of Monte Carlo chains *N*_chains_, with *m* = 1 fixed. In addition to *m*, the plateau level is also determined by the number of Monte Carlo chains, *N*_chains_. Decreasing *N*_chains_ can be seen as increasing the effective strength of regularization, helping to mitigate the overestimation of couplings. For sufficiently low values of *N*_chains_ (∼ 100), inference trajectories tend to slow down near the true value. At the same time, we observe significant variability between runs with identical *N*_chains_, because we use the stochasticity inherent to MCMC to navigate the optimization landscape.

**Figure S4:**
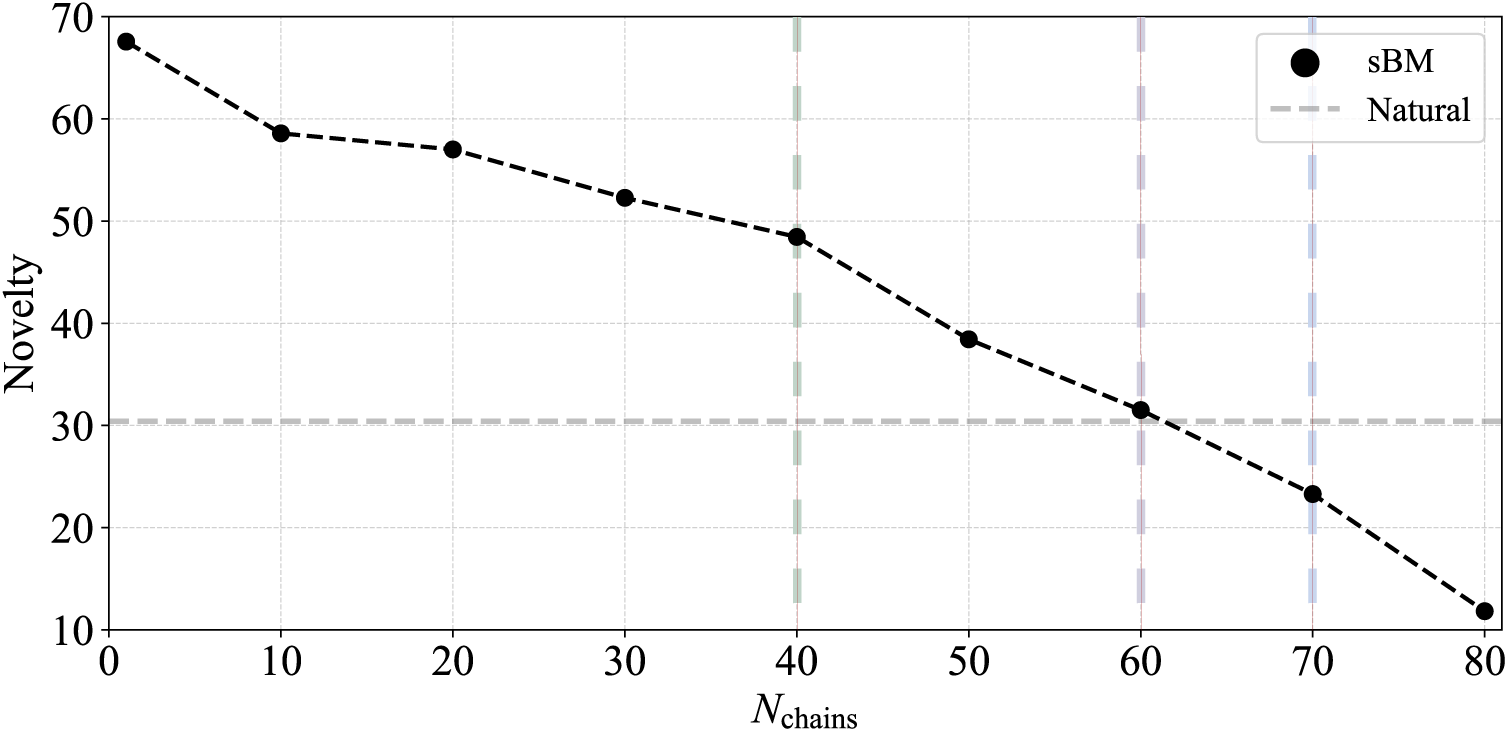
CM family: Novelty as a function of *N*_chains_ for sBM models. Novelty scores of sBM models inferred using different values of *N*_chains_. The horizontal dashed line indicates the novelty measured in the natural chorismate mutase MSA. The three vertical dashed lines indicate the values of *N*_chains_ corresponding to the three sBM models selected for experimental characterization. Increasing *N*_chains_ reduces the novelty of the generated sequences, illustrating how this parameter controls the regularization strength.

**Figure S5:**
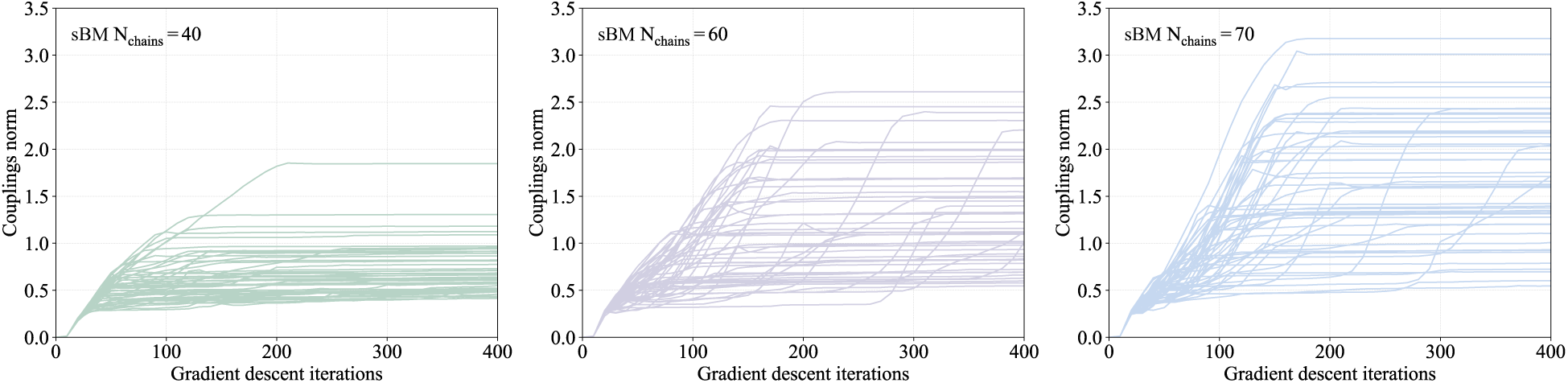
Chorismate Mutase family: couplings as a function of gradient descent iterations for sBM inferences. Average Frobenius norm of the couplings as a function of gradient descent iterations over 50 independent sBM inferences with *N*_chains_ = 40 (left), *N*_chains_ = 60 (middle), and *N*_chains_ = 70 (right). The variability observed across independent inferences highlights the stochastic nature of the sBM optimization procedure and motivates averaging model parameters over multiple runs. In particular, models used to assess generative performance on real data are obtained by averaging over 50 independent inferences.

**Figure S6:**
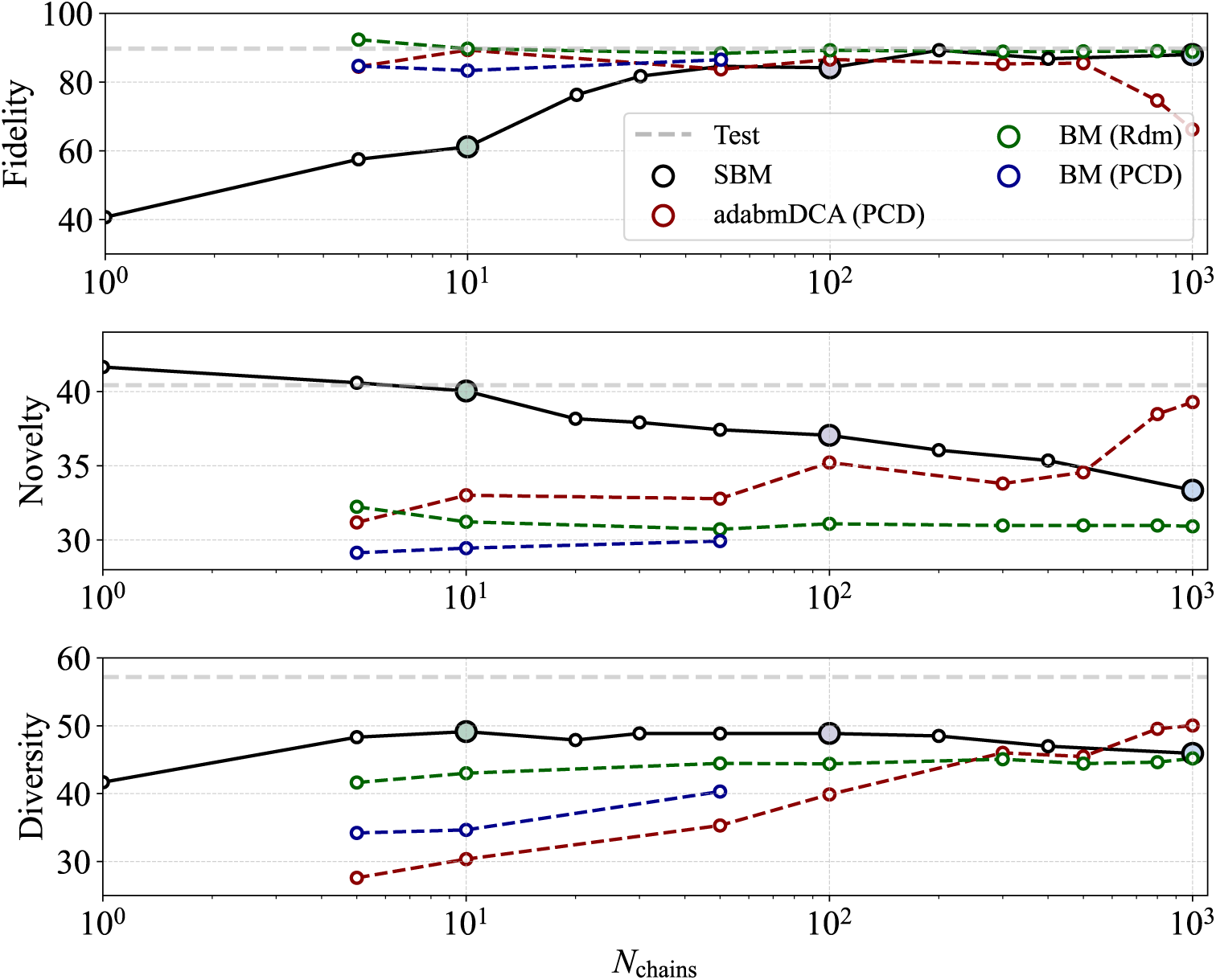
Generative performance of BM, sBM and adabmDCA [38] on synthetic data. Fidelity, novelty, and diversity scores of sequences generated by models inferred using different values of *N*_chains_. sBM models are shown in black. adabmDCA models, which use persistent contrastive divergence (PCD), are shown in red. To isolate the effect of PCD, BM models were inferred either by randomly initializing the Monte Carlo chains at each optimization step (BM (Rdm), green) or by reusing the final chain states from the previous optimization step (BM (PCD), blue). At low *N*_chains_, PCD substantially reduces the novelty and diversity of the generated sequences, showing that the effect of reducing *N*_chains_ depends on the Monte Carlo sampling procedure used during inference. Horizontal dashed lines indicate the corresponding scores of the teacher model.

**Figure S7:**
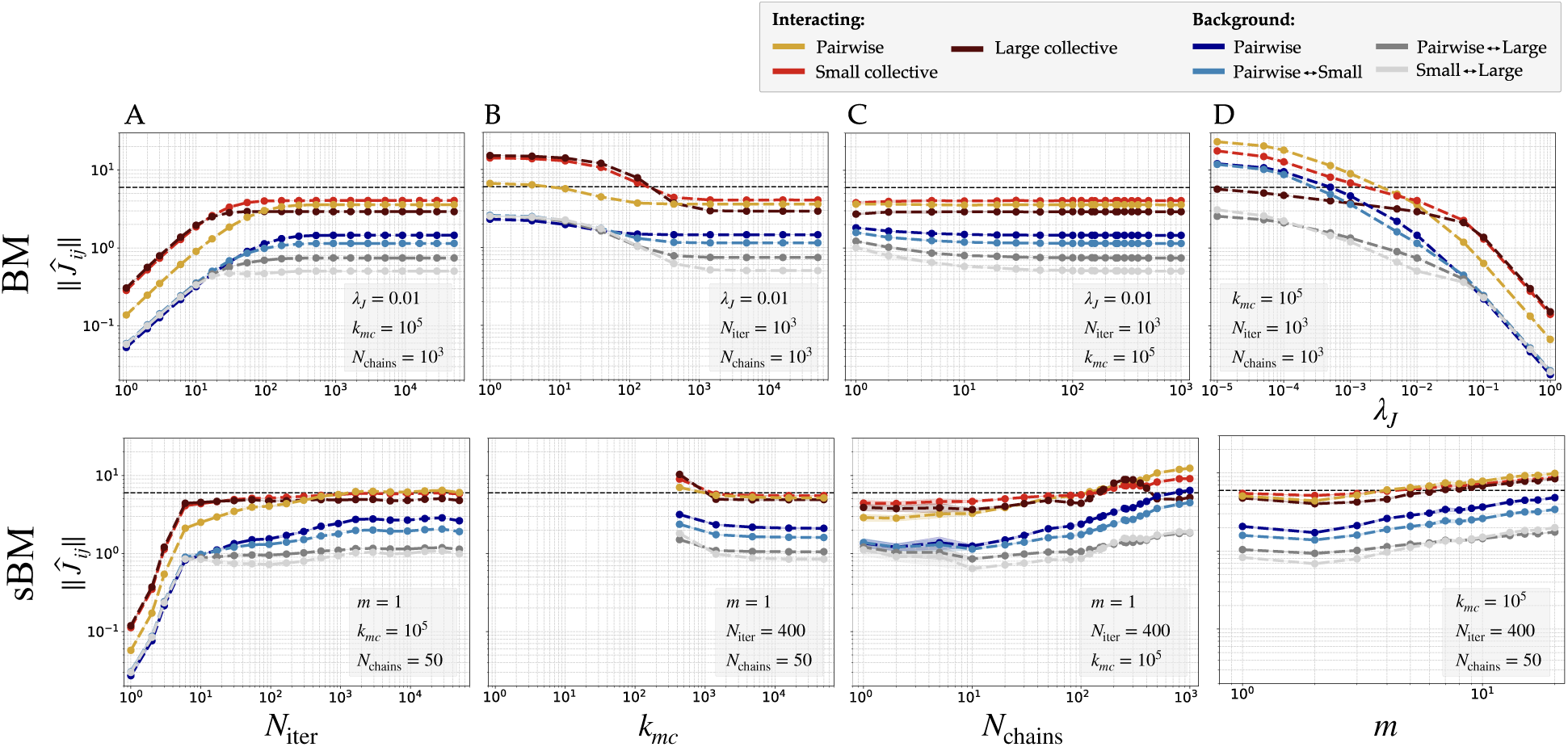
Comparison of BM and sBM inferences on the synthetic dataset from the mathematical model across hyperparameter choices. We evaluate the impact of various hyperparameters on the magnitude of inferred couplings ∥*Ĵ_ij_* ∥ for the different types of interacting couplings of the teacher model. The first row corresponds to results with the sBM, while the second row shows results with the BM. Coupling magnitudes are computed as Frobenius norms over amino acids of the coupling matrices, averaged over couplings belonging to the same category. The true coupling amplitude is shown as a dashed black line. Colors distinguish between true interacting couplings (pairwise: yellow, small collective: red, large collective: dark red) and background couplings (non-interacting) of different types (see legend and Fig. 2). Each column explores a different hyperparameter: **A.** Number of training iterations *N*_iter_. **B.** Number of MCMC steps *k*_mc_. **C.** Number of independent MCMC chains *N*_chains_. **D.** Regularization strength *λ_J_* for BM. For sBM, panel **D** instead varies the memory parameter *m* used in L-BFGS.

**Figure S8:**
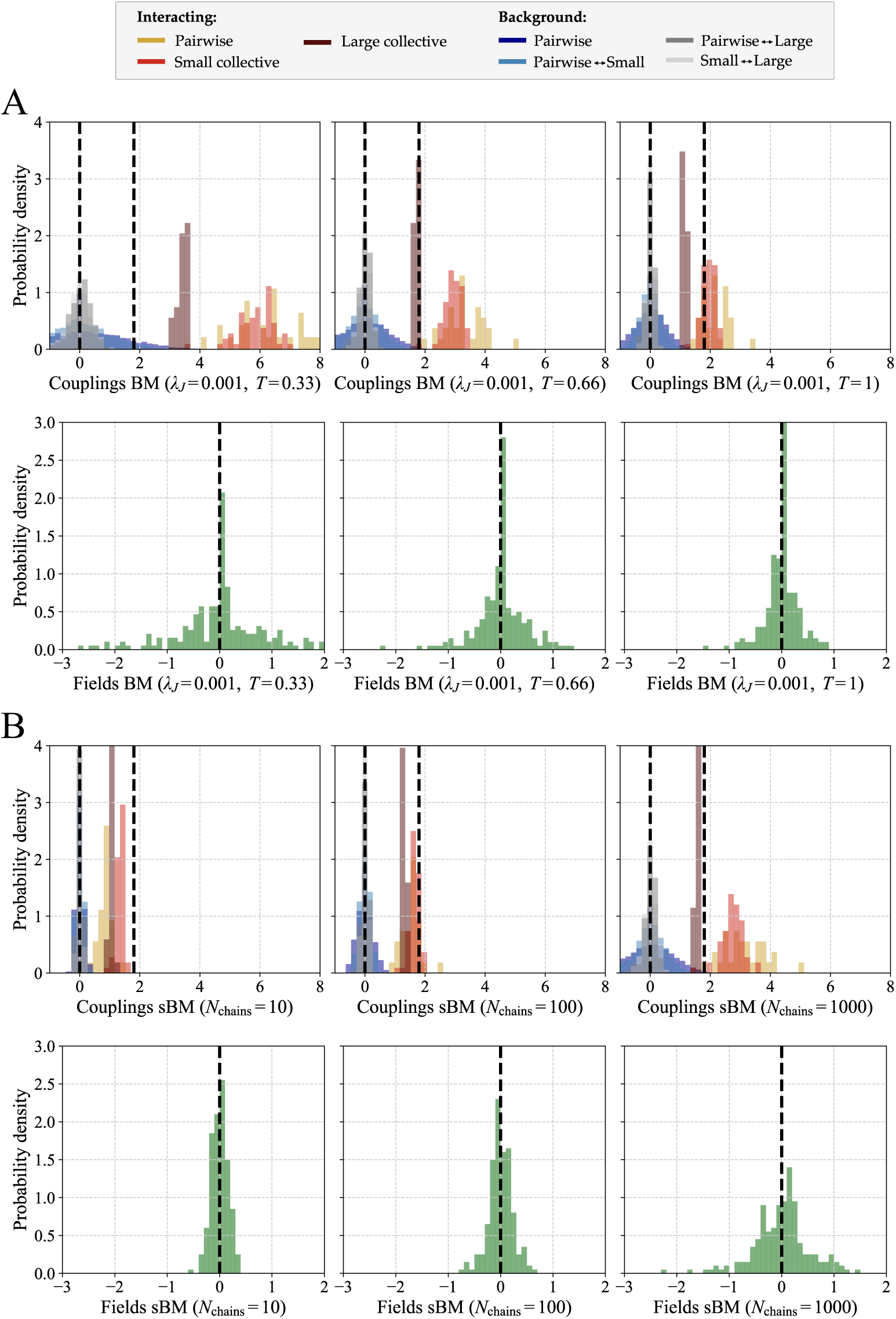
Distribution of inferred parameters in the context of synthetic data. Histograms of the inferred couplings (top row) and fields (bottom row). Couplings are colored according to their type in the teacher model (see legend). The vertical dashed lines indicate the corresponding ground-truth values of the interacting couplings and fields; background couplings and fields have a true value of zero. **A.** Parameters inferred with BM using an L2 regularization strength *λ_J_* = 0.001. From left to right, the three columns correspond to sampling temperatures *T* = 0.33, 0.66, and 1. **B.** Parameters inferred with sBM. From left to right, the three columns correspond to *N*_chains_ = 10, 100, and 1000.

**Figure S9:**
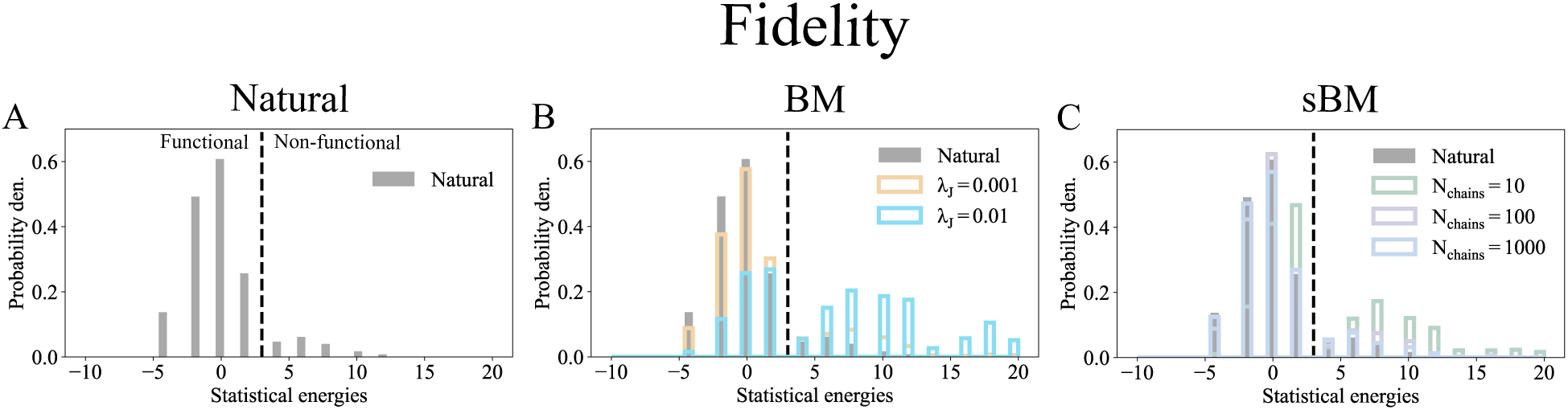
Synthetic data: fidelity in BM and sBM inferences. Distribution of the statistical energies of sequences generated with the teacher model (natural sequences), and with inferred models (artificial sequences). The statistical energies are calculated using the teacher model (see Fig. 1). **A.** Statistical energies of natural sequences. The functionality threshold is indicated by the vertical dotted line. **B.** Statistical energies of artificial sequences generated with BM inferences inferred with *λ* = 0.001 (orange) and *λ* = 0.01 (blue). As the regularization strength increases, the fraction of non-functional sequences also increases. **C.** Statistical energies of artificial sequences generated with sBM inferences inferred with *N*_chains_ = 10 (green), *N*_chains_ = 100 (purple) and *N*_chains_ = 1000 (blue). As the regularization strength increases, the fraction of non-functional sequences also increases.

**Figure S10:**
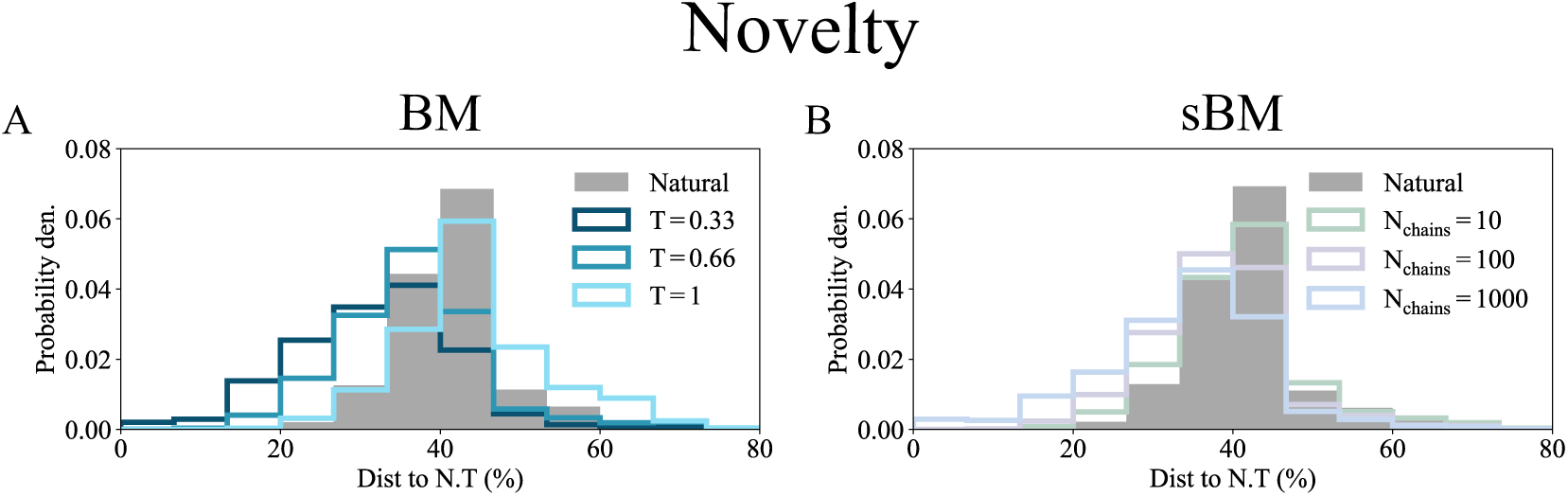
Synthetic data: novelty in BM and sBM inferences. **A.** Distribution of sequence identity to the nearest training sequence for a BM inference (*λ* = 0.01) at different sampling temperatures *T* = {0.33, 0.66, 1}. Lower temperatures produce sequences with higher similarity to the training data. **B.** Same analysis for sBM inferences with varying numbers of chains (*N*_chains_). Increasing *N*_chains_, i.e. reducing the regularization strength, results in sequences with higher similarity to the training data.

**Figure S11:**
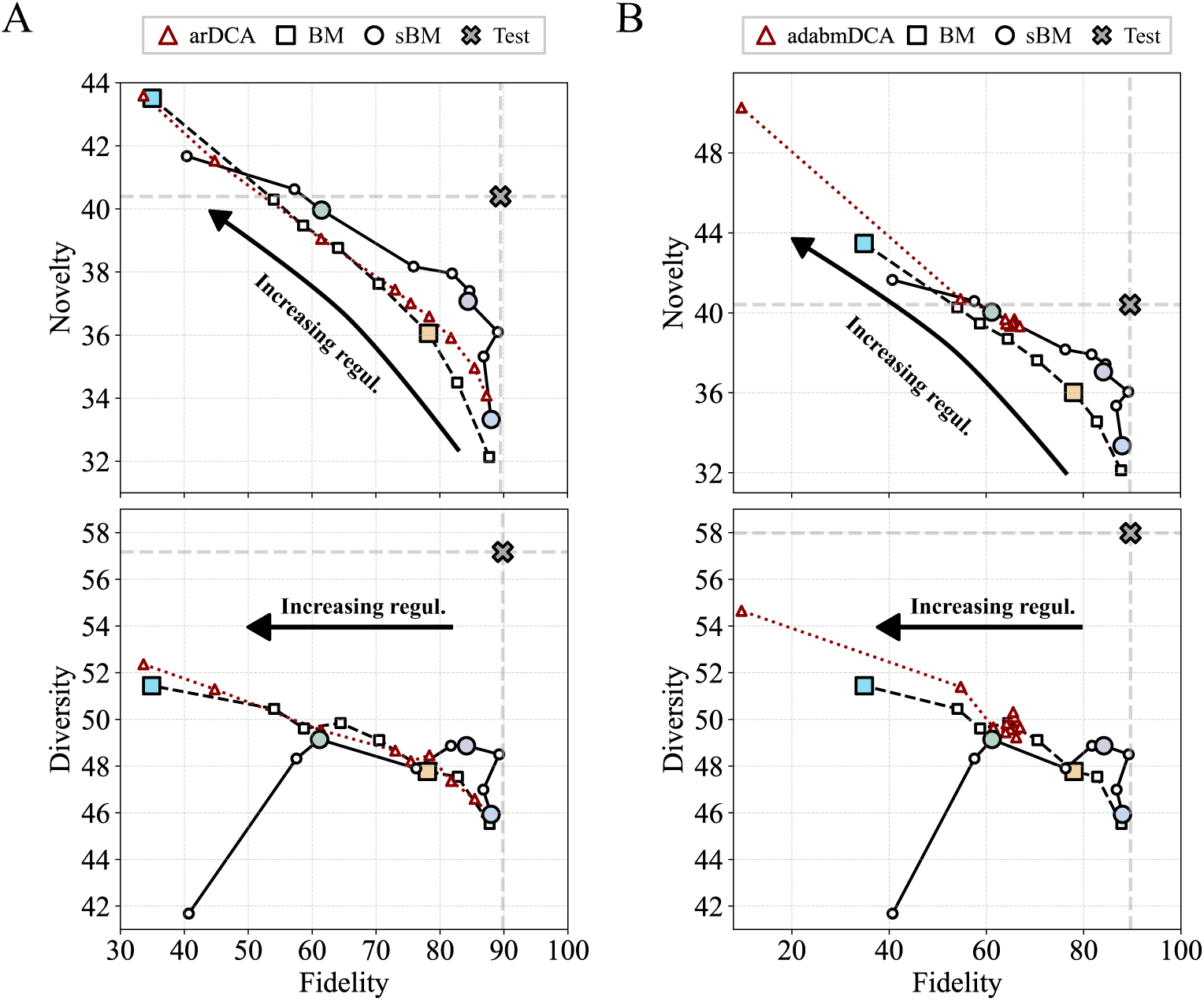
Generative performance of BM, sBM, arDCA [18], and adabmDCA [38] on synthetic data. Fidelity, novelty, and diversity scores (see Supplementary Methods) of sequences generated by models inferred with BM, sBM, arDCA, and adabmDCA. **A.** Comparison of sBM and BM with arDCA. The sBM models were inferred using different values of *N*_chains_ (from 1 to 1000, with *N*_chains_ = 1 corresponding to the strongest regularization), while the BM and arDCA models (in red) were inferred using different L2 regularization strengths (from 10*^−^*^5^ to 10*^−^*^2^). **B.** Comparison of sBM and BM with adabmDCA. The sBM and BM results are identical to those shown in panel A. The red symbols correspond to adabmDCA models inferred without L2 regularization, using different pseudocount values (from 10*^−^*^5^ to 5 × 10*^−^*^1^). All other adabmDCA hyperparameters were kept at their default values, except for *N*_chains_, which was set to 1000.

**Figure S12:**
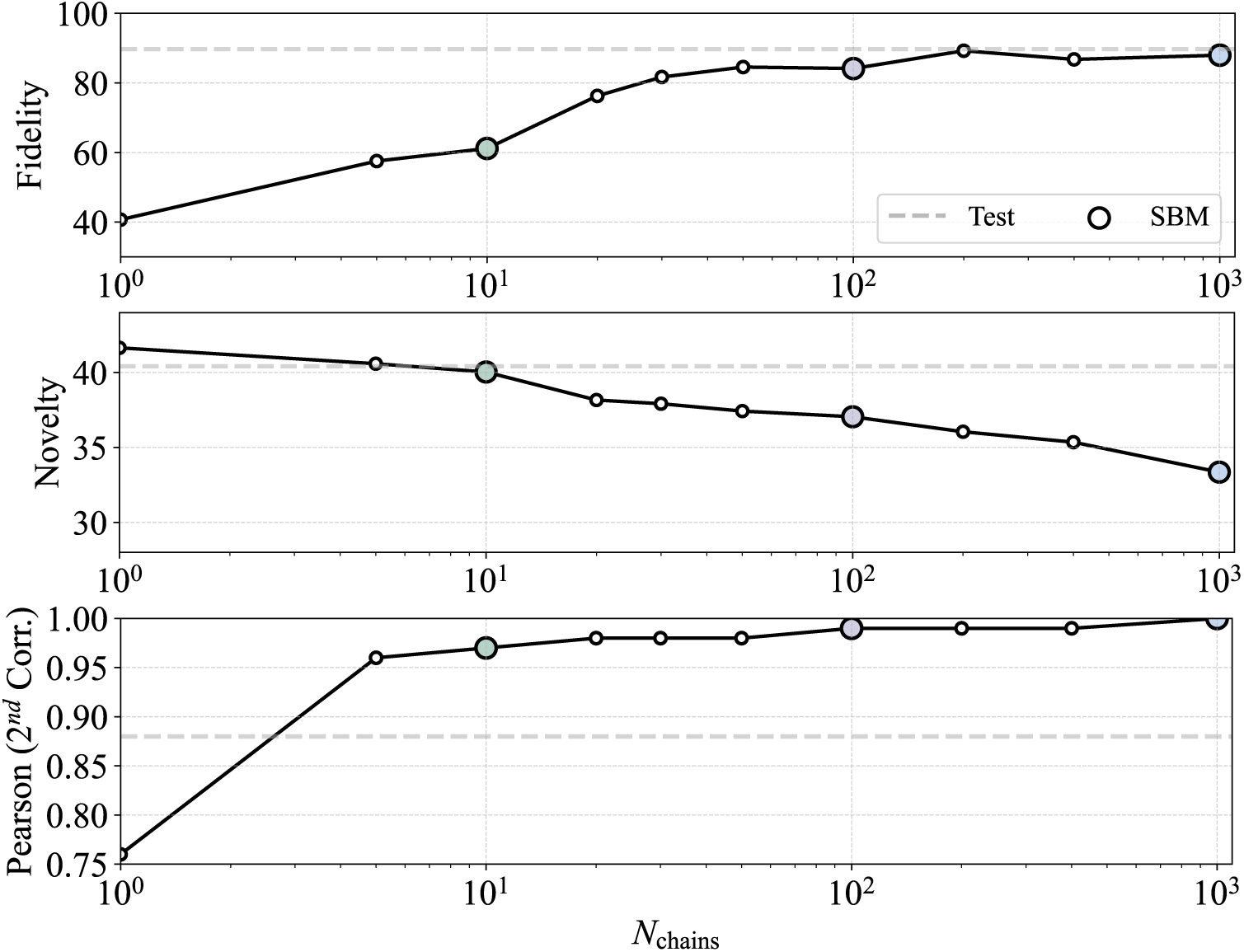
Comparison of statistical fit and generative performance in the context of synthetic data. Evolution of the fidelity (top), novelty (middle), and Pearson correlation between the empirical and generated second-order statistics (bottom) as a function of *N*_chains_ for sBM. The horizontal dashed lines indicate the corresponding values obtained for the teacher model. As the inferred model approaches a perfect reproduction of the empirical second-order statistics (Pearson correlation approaching one), the novelty of the generated sequences decreases markedly. Conversely, models that reproduce the empirical statistics slightly less accurately achieve a substantially better fidelity–novelty trade-off.

**Figure S13:**
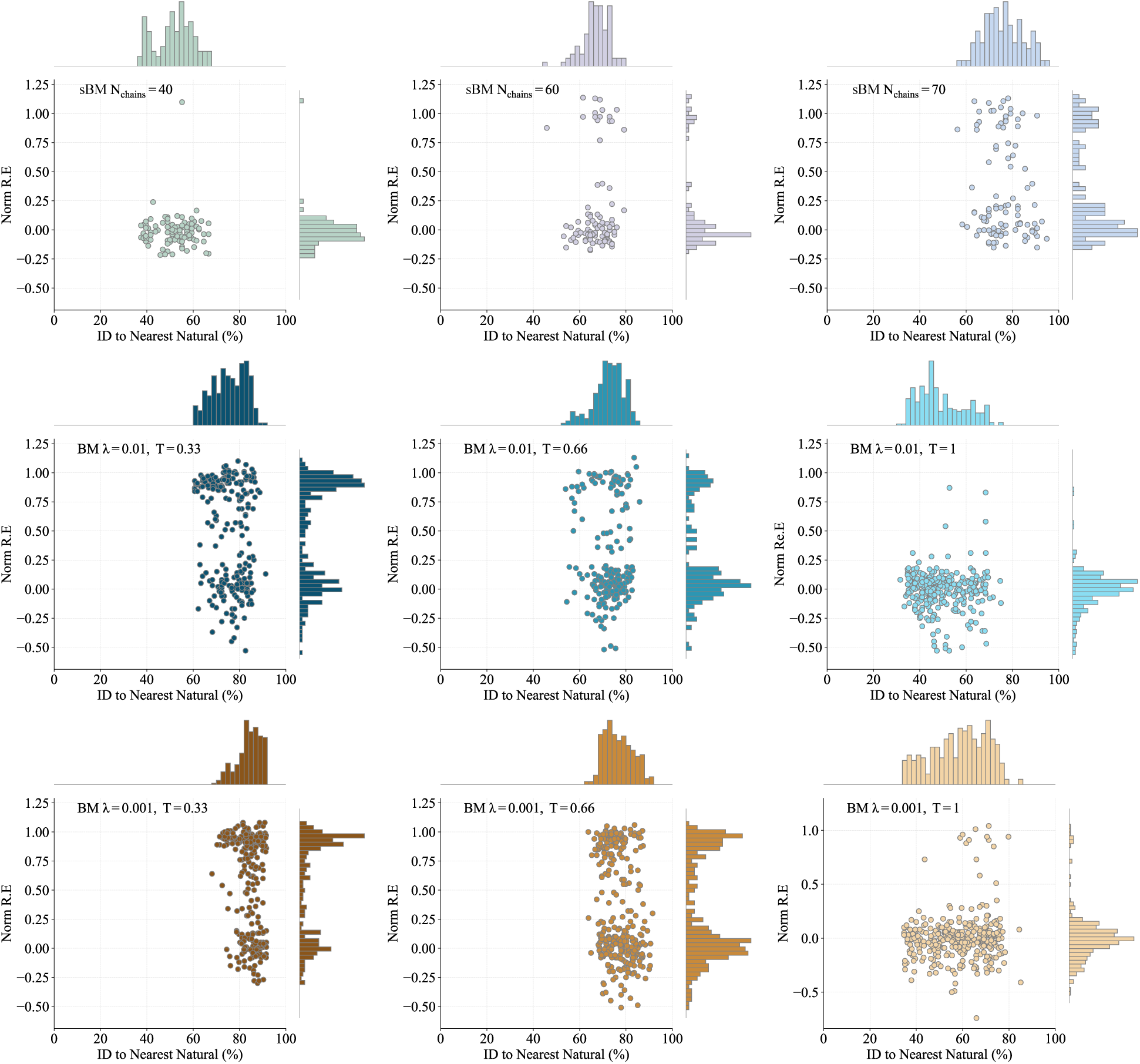
Chorismate mutase family: novelty and fidelity in BM and sBM inferences. Scatter plots showing the relative enrichment measured experimentally as a function of identity to the nearest natural sequence, for artificial sequences generated by BM and sBM inferences. **Upper row.** Artificial sequences generated with sBM inference using *N*_chains_ = 40 (left), *N*_chains_ = 60 (middle) and *N*_chains_ = 70 (right). **Middle row.** Artificial sequences generated with BM inference using *λ* = 0.001 and sampling using *T* = 0.33 (left), *T* = 0.66 (middle) and *T* = 1 (right). These plots were made using Russ *et al.* data [11]. **Lower row.** Artificial sequences generated with BM inference using *λ* = 0.01 and sampled using *T* = 0.33 (left), *T* = 0.66 (middle) and *T* = 1 (right). These plots were made using Russ *et al.* data [11].

**Figure S14:**
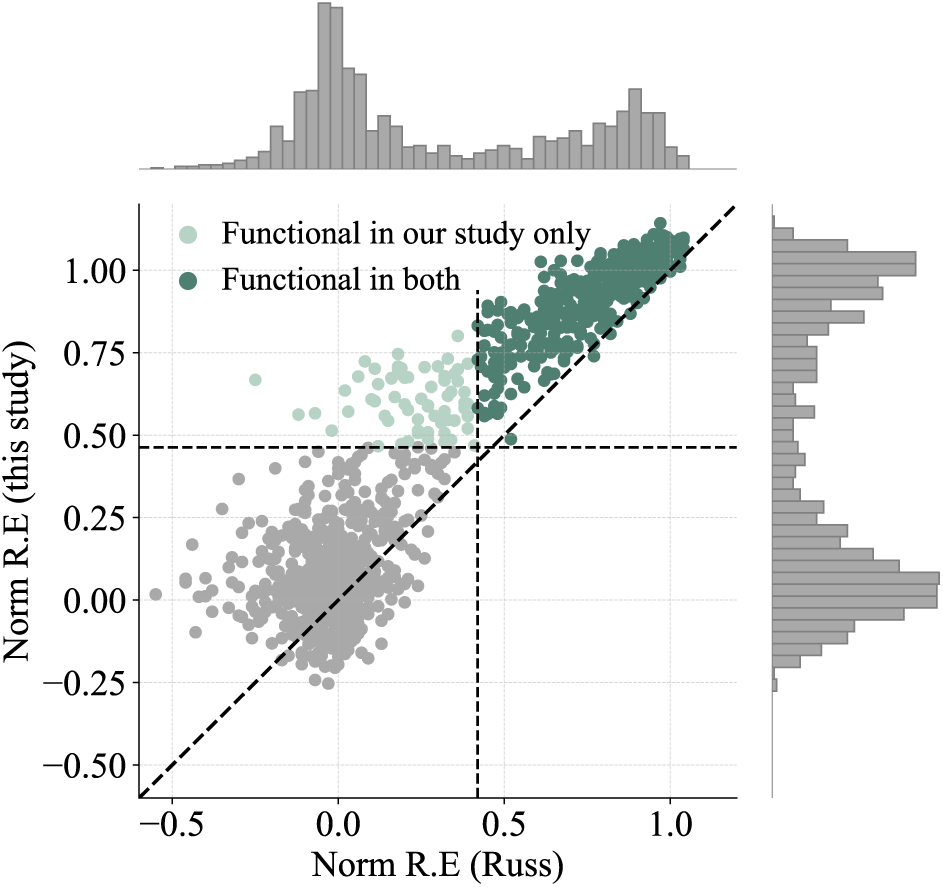
Chorismate mutase family: comparison of normalized relative enrichment (R.E.) values for natural sequences between this study and Russ *et al.* [11]. Each point represents a natural sequence. The normalized relative enrichment obtained in this study is plotted against the corresponding value reported by Russ *et al.*. Dashed lines indicate functional thresholds used in each dataset. Dark green points denote sequences identified as functional in both studies, while light green points are considered to be functional only in this study.

**Figure S15:**
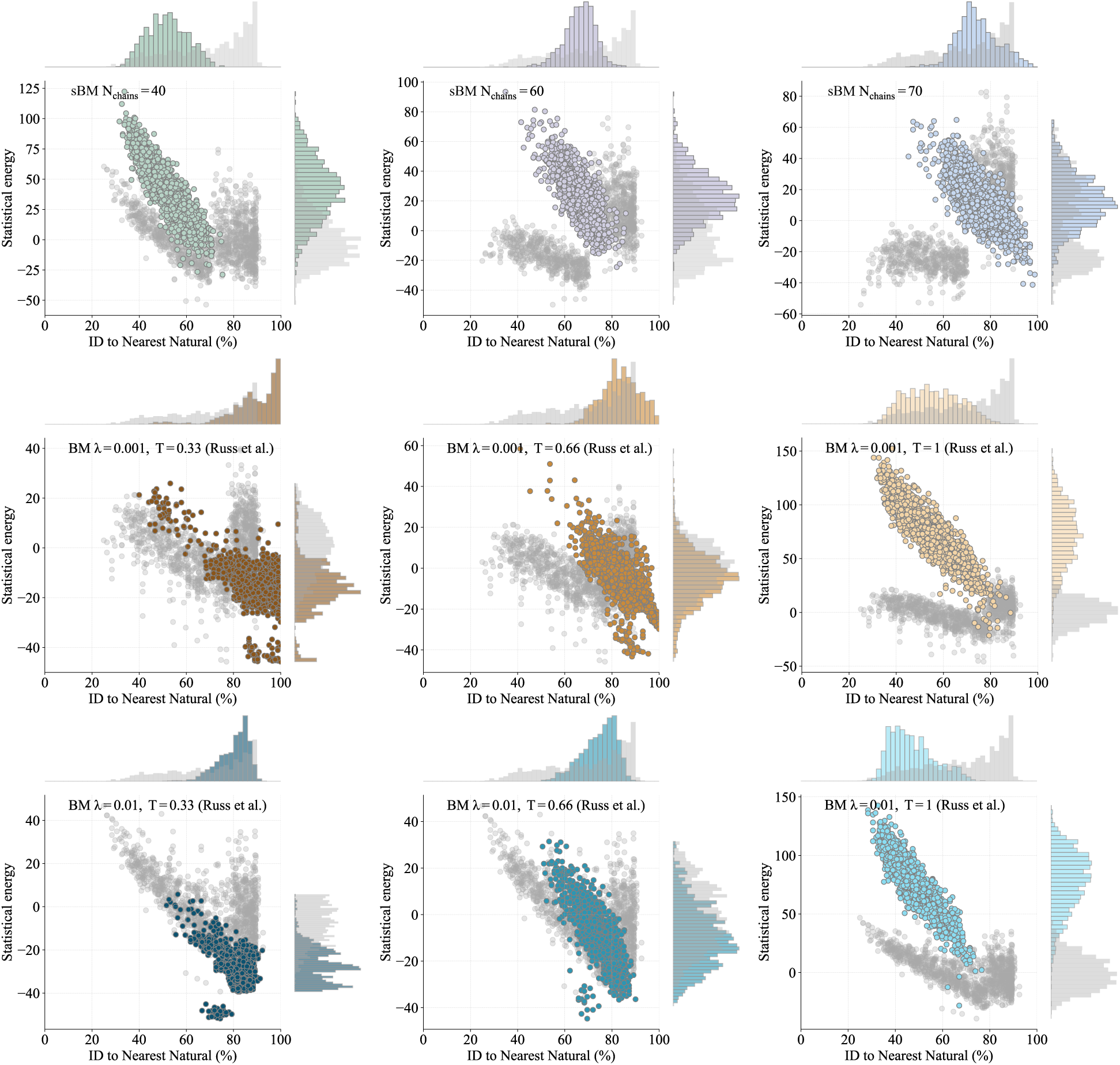
Chorismate mutase family: Scatter plots showing the statistical energies of each artificial sequence against its identity to the nearest natural sequence. **Upper row.** Artificial sequences generated by sBM inference with *N*_chains_ = 40 (left), *N*_chains_ = 60 (middle) and *N*_chains_ = 70 (right). **Middle row.** Artificial sequences generated by BM inference with *λ* = 0.001 and sampled with *T* = 0.33 (left), *T* = 0.66 (middle) and *T* = 1 (right). These plots are made using Russ *et al.* data [11]. **Lower row.** Artificial sequences generated by BM inference with *λ* = 0.01 and sampled with *T* = 0.33 (left), *T* = 0.66 (middle) and *T* = 1 (right). These plots are made using Russ *et al.* data [11].

**Figure S16:**
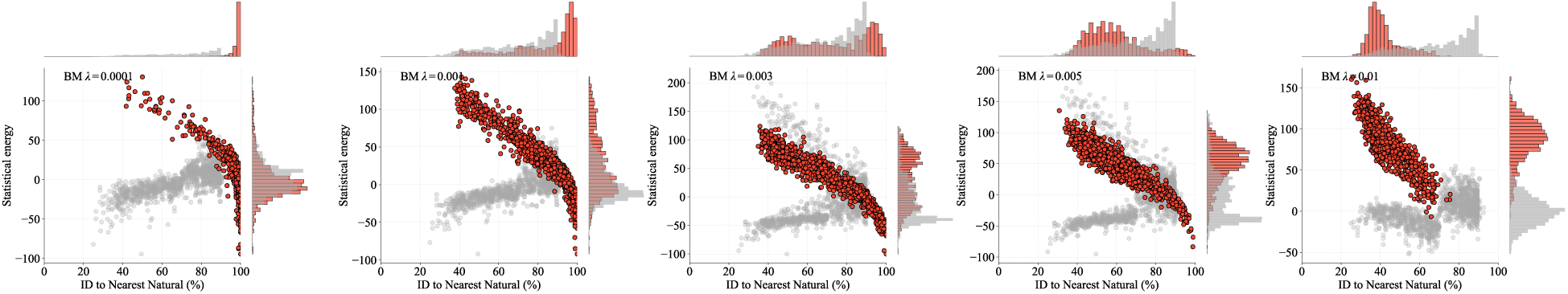
Chorismate mutase family: models inferred with BM cannot simultaneously match natural energies and generate novel sequences. Each panel reports statistics for sequences sampled at *T* = 1 from a BM trained on the chorismate mutase family with different *L*_2_ regularization strengths: *λ* = 0.001, 0.003, 0.005, 0.01 (from left to right). Percentage identity (ID) to the closest natural sequence against statistical energies for natural (light grey), and BM-generated sequences (red). Lowering the regularization (smaller *λ*) drives the model to sample artificial sequences with statistical energies similar to those of the training alignment, but at the cost of producing sequences that are almost all identical to natural ones. Conversely, stronger regularization (larger *λ*) yields more diverse sequences but shifts their energy distribution away from that of the training data.

**Figure S17:**
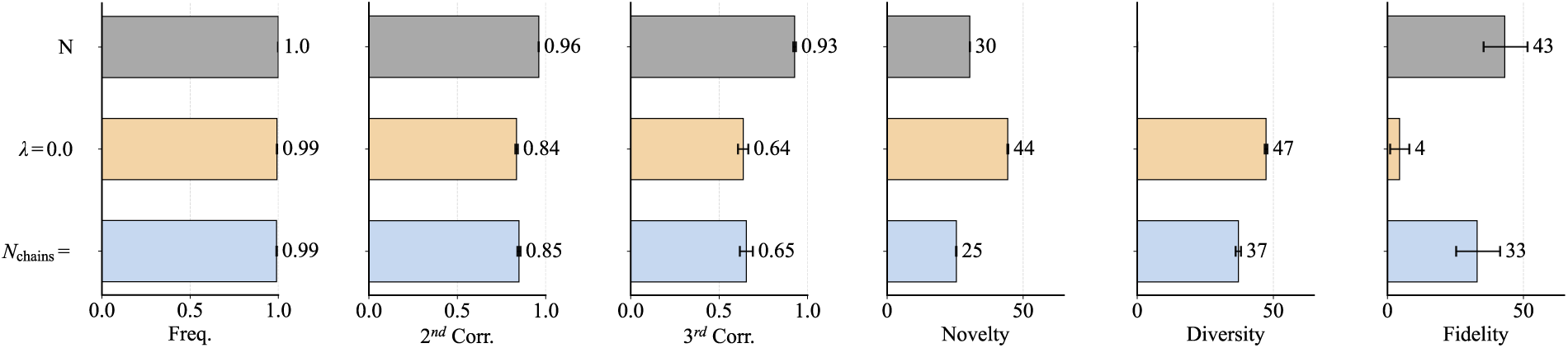
CM family: Comparison of statistical fit and generative performance. Comparison of the BM model of Russ *et al.* [11] (*λ* = 0.001) and the sBM model inferred with *N*_chains_ = 70. Statistical performance is quantified by the Pearson correlation between the empirical first-, second-, and third-order statistics of the natural MSA and those computed from sequences generated with each model. Although the two models reproduce the natural statistics with comparable accuracy, they differ substantially in novelty, diversity, and fidelity, demonstrating that matching empirical statistics alone is not sufficient to assess the generative performance of protein sequence models.

**Figure S18:**
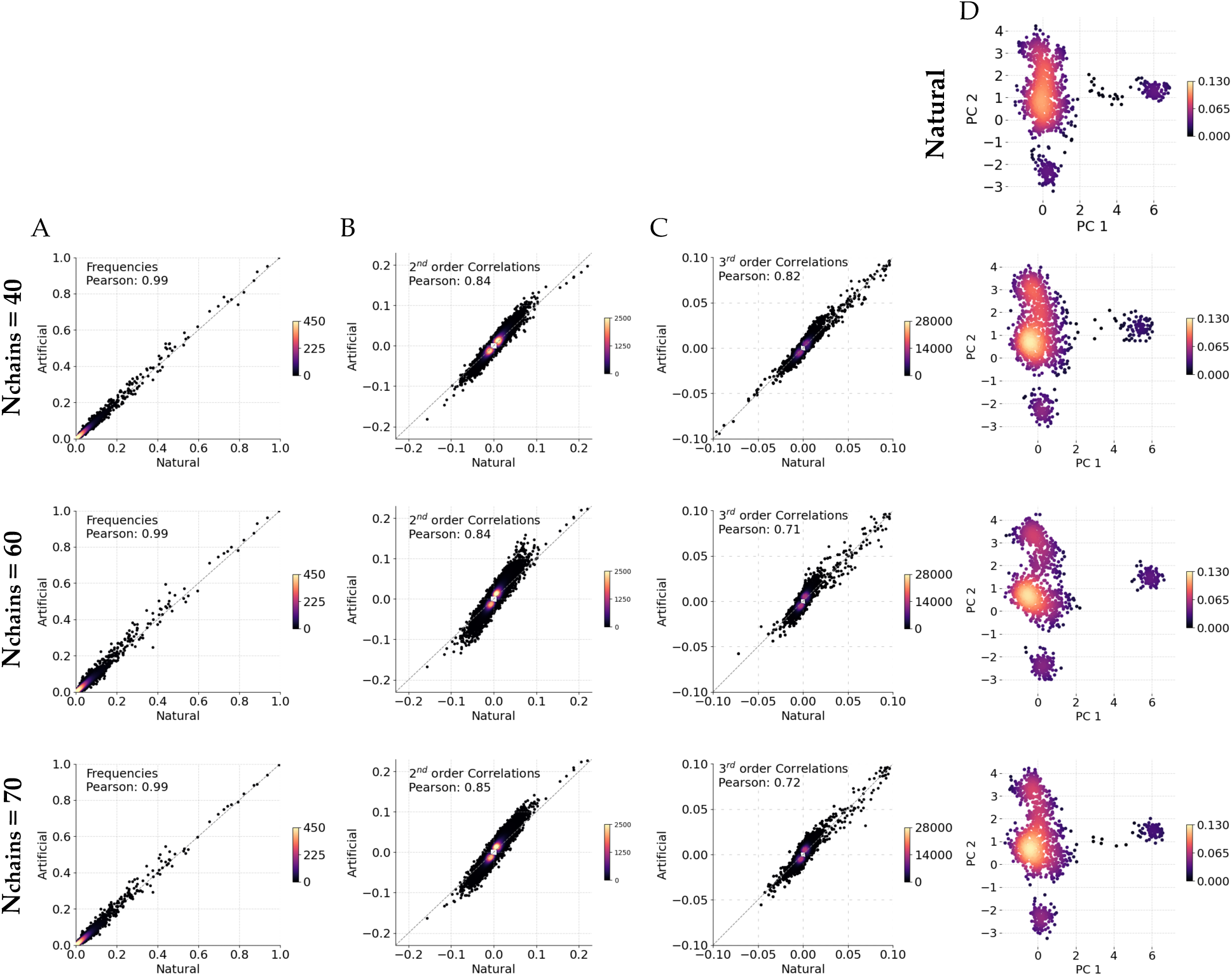
Chorismate mutase family: comparison of statistical properties between natural sequences and artificial sequences generated by sBM inferences. We compare statistical properties computed from artificial sequences generated by sBM inference with different regularization strengths to those computed from the natural sequence alignment. Rows correspond to *N*_chains_ = 40 (top), 60 (middle), and 70 (bottom). **A**. Comparison of single-site amino-acid frequencies between natural sequences and artificial sequences. Pearson correlation coefficients are indicated and point density is encoded by color. **B**. Same comparison as in panel A, but for pairwise (second-order) correlations. **C**. Same comparison as in panel A, but for third-order correlations. **D**. Projection of sequences onto the first two principal components (PC1 and PC2) computed from the natural sequence alignment. Each point represents a sequence, colored by local density. The top panel shows the corresponding projection of natural sequences, used as a reference.

